# Engineering and Evolution of Methanol Assimilation in *Saccharomyces cerevisiae*

**DOI:** 10.1101/717942

**Authors:** Monica I. Espinosa, Ricardo A. Gonzalez-Garcia, Kaspar Valgepea, Manuel Plan, Colin Scott, Isak S. Pretorius, Esteban Marcellin, Ian T. Paulsen, Thomas C. Williams

## Abstract

Microbial fermentation for chemical production is becoming more broadly adopted as an alternative to petrochemical refining. Fermentation typically relies on sugar as a feedstock, however, one-carbon compounds like methanol are an attractive alternative as they can be derived from organic waste and natural gas. This study focused on engineering methanol assimilation in the yeast *Saccharomyces cerevisiae.* Three methanol assimilation pathways were engineered and tested: a synthetic xylulose monophosphate (XuMP), a ‘hybrid’ methanol dehydrogenase-XuMP, and a bacterial ribulose monophosphate (RuMP) pathway, with the latter identified as the most effective at assimilating methanol. Additionally, ^13^C-methanol tracer analysis uncovered a native capacity for methanol assimilation in *S. cerevisiae*, which was optimized using Adaptive Laboratory Evolution. Three independent lineages selected in liquid methanol-yeast extract medium evolved premature stop codons in *YGR067C*, which encodes an uncharacterised protein that has a predicted DNA-binding domain with homology to the *ADR1* transcriptional regulator. Adr1p regulates genes involved in ethanol metabolism and peroxisomal proliferation, suggesting *YGR067C* has a related function. When one of the evolved *YGR067C* mutations was reverse engineered into the parental CEN.PK113-5D strain, there were up to 5-fold increases in ^13^C-labelling of intracellular metabolites from ^13^C-labelled methanol when 0.1 % yeast extract was a co-substrate, and a 44 % increase in final biomass. Transcriptomics and proteomics revealed that the reconstructed *YGR067C* mutation results in down-regulation of genes in the TCA cycle, glyoxylate cycle, and gluconeogenesis, which would normally be up-regulated during growth on a non-fermentable carbon source. Combining the synthetic RuMP and XuMP pathways with the reconstructed Ygr067cp truncation led to further improvements in growth. These results identify a latent methylotrophic metabolism in *S. cerevisiae* and pave the way for further development of native and synthetic one-carbon assimilation pathways in this model eukaryote.

## Introduction

Our current dependence on fossil fuels is not sustainable due to their finite reserves and the negative environmental impacts caused by their use. By-products from fossil fuel combustion include a myriad of toxic air pollutants and CO_2_, which is the main anthropogenic contributor to climate change. These complex environmental problems call for a global effort to move towards a bio-economy in which microbial metabolism is used for the conversion of renewable materials into useful products ^1^. Typically, sugars derived from sugarcane or corn are used as feedstocks for the production of fuels and chemicals using biological fermentation. However, sugar production is costly and requires arable land that competes with other land uses, such as food production. It is estimated that the sugar cost represents up to 70 % of the total production costs ^2^, making many biotechnological processes uncompetitive with petrochemical processes.

Growing crops for feedstock also requires water and fertilizer, which could be instead allocated to grow food to meet the demand of an ever-rising population. Alternatively, lignocellulosic non-food biomass could be used for obtaining sugars, but the process is limited by the recalcitrance of the feedstock and difficulties utilising the large lignin fraction of biomass. One-carbon (C1) metabolites are abundant and inexpensive, and can be obtained from natural gas or waste resources such as agricultural, municipal or industrial waste, or natural gas ^3, 4^. Methanol in-particular is becoming an attractive feedstock due to its abundance and liquid state, which makes it more compatible with existing fermentation, storage, and transportation infrastructure ^3, 5^. Methanol can also be obtained inexpensively from methane and carbon dioxide stemming from industrial waste streams ^6^. Engineering microorganisms to grow on and convert C1 compounds such as methanol into food, fuels, and chemicals has therefore become a major goal in the fields of synthetic biology and metabolic engineering ^7, 8^.

Methylotrophic metabolism enables energy and biomass generation from methanol, and is present in bacterial, archaeal, and yeast species. Recent attempts have been made to produce valuable metabolites using methylotrophs. For example, *Methylobacterium extorquens* AM1 has been engineered to produce 3-hydroxypropionic acid from methanol ^9^. The methylotrophic yeast *Pichia pastoris* (renamed *Komagataella phaffii* ^10^) is currently used industrially for production of recombinant proteins ^11^. Recently, *P. pastoris* has also been used for production of metabolites including astaxanthin and isobutanol ^12, 13^. However, relative to model organisms, native methylotrophs lack the genetic tools and depth of characterisation necessary for the successful metabolic engineering of high-yield and heterologous pathways. An attractive alternative to enable the use of methanol as feedstock is to engineer synthetic methylotrophy into industrially-robust and well-characterised microorganisms. Model organisms are not only easier to genetically manipulate and optimise, but also provide the opportunity to engineer non-native central carbon metabolism that has greater genetic and metabolic plasticity. Recent approaches have focused on engineering synthetic methylotrophy in bacteria such as *Escherichia coli* and *Corynebacterium glutamicum* ^14–21^. Incorporation of ^13^C-methanol into central carbon metabolites and specialty products has been demonstrated in both species; however, growth on methanol still requires yeast extract or additional carbon sources like xylose.

The yeast *Saccharomyces cerevisiae* has potential as a host for the engineering of synthetic methylotrophy as it has distinct advantages over organisms such as *E. coli* for use in industrial fermentation. For example, *S. cerevisiae* can correctly express, fold, and post-translationally modify eukaryotic proteins, is not susceptible to phage contamination, has high tolerance to low pH concentrations and solvents like methanol, and has organelles that can be co-opted for localization of specialized metabolism ^22–24^. The engineering of synthetic methylotrophic metabolism in *S. cerevisiae* could therefore provide several benefits for methanol-fed bioprocesses. Here, we designed, built, and tested three different methanol-assimilation pathways in *S. cerevisiae* and identified for the first time the native capacity of *S. cerevisiae* to assimilate methanol into central carbon metabolism. Using methanol-dependent growth assays, ^13^C-methanol tracer studies, and Adaptive Laboratory Evolution (ALE), we characterised synthetic methanol assimilation pathways and optimised native methanol assimilation in *S. cerevisiae*.

## Results

### Benchmarking synthetic pathways for methanol assimilation potential

After evaluating the different methanol assimilation pathways that have been previously described in nature, three metabolic pathways were designed for engineering synthetic methylotrophy in *S. cerevisiae*. The first pathway design (coloured red in Fig. 1A) was based on the xylulose monophosphate (XuMP) pathway in the methylotrophic yeast *P. pastoris.* Here, methanol assimilation genes are targeted to the peroxisome by a type 1 peroxisome targeting signal (PTS1). An FAD-dependent alcohol oxidase (Aox1p) oxidises methanol to formaldehyde and hydrogen peroxide. Hydrogen peroxide is detoxified by a catalase (Cat1p) in the peroxisome while formaldehyde is converted to glyceraldehyde-3-phosphate by dihydroxyacetone synthase (Das1p), which uses xylulose-5-phosphate as a co-substrate. Glyceraldehyde-3-phosphate can then be incorporated into central carbon metabolism to produce biomass. The second pathway was termed a ‘hybrid’ because it links the *P. pastoris* XuMP pathway with an NAD^+^-dependent methanol dehydrogenase (Mdh) from *Bacillus stearothermophilus*, which oxidises methanol to formaldehyde but does not produce hydrogen peroxide. Formaldehyde is then converted to glyceraldehyde-3-phosphate by a cytosolically localised Das1p from *P. pastoris*. Utilising Mdh has the advantage of not requiring oxygen for the reaction and not producing hydrogen peroxide, unlike Aox1p, and generating energy in the form of NADH. The third pathway was based on the bacterial ribulose monophosphate (RuMP) pathway where Mdh oxidises methanol to formaldehyde while hexulose-6-phosphate synthase (Hps) uses ribulose-5-phosphate as a co-substrate to convert formaldehyde to hexulose-6-phosphate, which is then converted to fructose-6-phosphate by 6-phospho-3-hexuloisomerase (Phi) from *Bacillus methanolicus*. Fructose-6-phosphate can then enter glycolysis to produce biomass precursors.

**Fig. 1.**
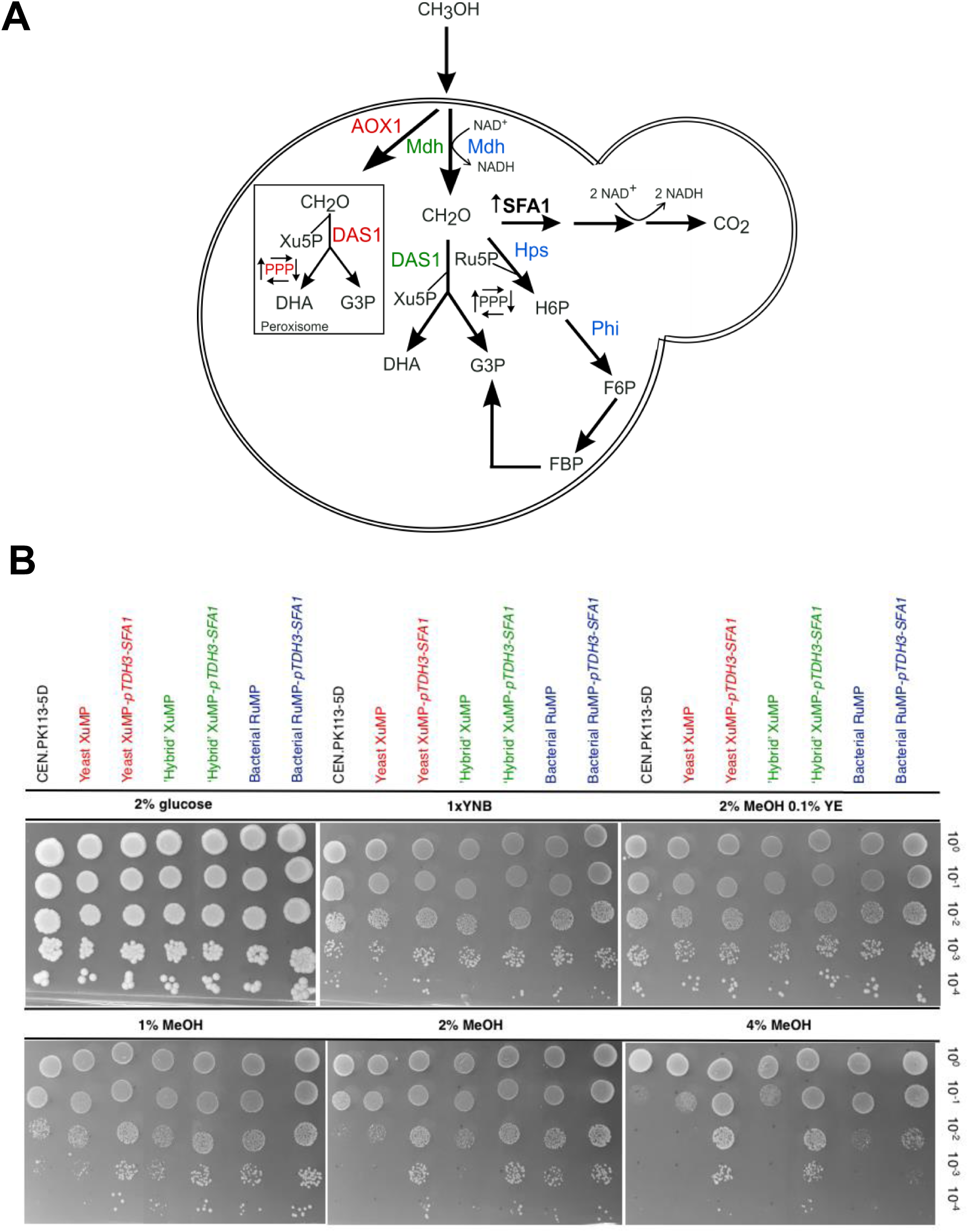
Methanol assimilation pathways engineered in *S. cerevisiae*. **A.** Three different metabolic pathways were engineered for methanol assimilation where methanol is oxidised to formaldehyde and later assimilated to glyceraldehyde-3-phosphate or fructose-6-phosphate. The first yeast XuMP pathway (red) is targeted to the peroxisome. The second ‘hybrid’ XuMP pathway (green) occurs in the cytosol. The third bacterial RuMP pathway (blue) is also targeted to the cytosol and converts formaldehyde to fructose-6-phosphate. Over-expression of the native formaldehyde detoxification enzyme Sfa1p is shown in bold. AOX1, alcohol oxidase 1; DAS1, dihydroxyacetone synthase; PPP, pentose phosphate pathway; Mdh, methanol dehydrogenase; Hps, hexulose-6-phosphate synthase; Phi, phospho-3-hexuloisomerase; SFA1, S-(hydroxymethyl) glutathione dehydrogenase; Xu5P, xylulose-5-phosphate; Ru5P, ribulose-5-phosphate. **B.** Growth assays of different methanol assimilation pathways with or without over-expression of the native formaldehyde detoxification pathway. Growth on increasing concentrations of methanol was tested in serial 10-fold dilutions for an empty vector control, the yeast XuMP (red), ‘hybrid’ XuMP (green), or bacterial RuMP (blue) strain with or without over-expression of *SFA1*. Yeast Nitrogen Base (YNB), Yeast Extract (YE), Methanol (MeOH). Images were taken after incubating at 30 °C for 5 days.

The three pathways, yeast XuMP, ‘hybrid’ XuMP, and bacterial RuMP, were expressed from low-copy centromeric plasmids in *S. cerevisiae* and tested by spotting serial 10-fold dilutions of the strains onto solid minimal Yeast Nitrogen Base (YNB) medium with 1 %, 2 %, or 4 % methanol alongside an empty vector control (CEN.PK113-5D; Fig. 1B). These spot assays are a high-throughput method for assaying growth, with colony size and spot-density comparisons used to identify growth differences. To determine if the growth differences were methanol-specific, strains were also spotted onto solid minimal YNB medium with 2 % glucose, YNB medium with no added carbon source (1 x YNB) or 2 % methanol with 0.1 % yeast extract (Fig. 1B). The yeast XuMP pathway (*P. pastoris AOX1* and *DAS1*) resulted in a subtle growth increase on 1 % methanol medium compared to the empty vector control, but this was not visible at 2 % methanol. The ‘hybrid’ XuMP pathway with bacterial *mdh* and *P. pastoris DAS1* expression had increased growth on 1 % and 2 % methanol relative to the control strain. The same was observed in the bacterial RuMP pathway with *mdh*, *hps*, and *phi*. Growth was still observed on 1 x YNB medium with no added carbon source, indicating there are medium components that enable the observed background growth. Strain-specific growth differences were far more pronounced on YNB-methanol media without yeast-extract.

### Enhancing native formaldehyde detoxification improves methanol-specific growth

Methylotrophy is a delicate balance between formaldehyde assimilation and dissimilation. Formaldehyde dissimilation to CO_2_ is the main source of energy for methylotrophs, as the TCA cycle is down-regulated when methanol is the sole carbon source ^25^. While formaldehyde assimilation *in S. cerevisiae* requires heterologous enzyme expression, a native pathway for the dissimilation of formaldehyde to CO_2_ and energy in the form of NADH already exists (Fig. 1A) ^26, 27^. This pathway is initiated by an NAD^+^ and glutathione-dependent formaldehyde dehydrogenase, Sfa1p, which has previously been shown to enhance formaldehyde tolerance when over-expressed in *S. cerevisiae* ^28, 29^. As formaldehyde accumulation in the cell can be toxic, and its dissimilation plays an important role in methylotrophs, we chose to over-express *SFA1* in *S. cerevisiae* alongside each of the three methanol assimilation pathways.

Growth of the yeast XuMP, ‘hybrid’ XuMP, or bacterial RuMP strains with or without over-expression of *SFA1* was tested on solid minimal YNB medium with increased concentrations of methanol or 2 % methanol with 0.1 % yeast extract, YNB medium with no added carbon source (1 x YNB), or 2 % glucose (Fig. 1B). To confirm methanol assimilation, growth on methanol would have to be visible independently of *SFA1* over-expression. For example, from Fig. 1B a higher capacity for growth can be observed on 1 and 2 % methanol for the ‘hybrid’ XuMP and bacterial RuMP strains. Enhanced formaldehyde dissimilation via *SFA1* over-expression had a further positive effect on the growth of these strains at 2 % and 4 % methanol (Fig. 1B). This distinct growth increase was most pronounced in the bacterial RuMP strain relative to both the empty vector control strain on the same media, and relative to the same engineered strains on 1 x YNB medium. This experiment indicated three critical things: (i) the RuMP and ‘hybrid’ pathways enabled methanol and formaldehyde assimilation in *S. cerevisiae*; (ii) the RuMP pathway is superior to the ‘hybrid’ for formaldehyde assimilation; and (iii) *SFA1* over-expression is an effective method for formaldehyde dissimilation, which is beneficial for growth on methanol.

Further modifications aimed at improving growth on methanol of the ‘hybrid’ XuMP and bacterial RuMP strains included targeting of *DAS1* to Ty1 sites for multiple genome insertion or including the second enzymatically active dihydroxyacetone synthase (Das2p) in *P. pastoris* in a new ‘hybrid’ XuMP *DAS1/DAS2* strain, as well as attempting to modulate flux through the pentose phosphate pathway by over-expressing *TKL1* or *TAL1* in either strain (Supplementary Fig. 1). Over-expression of these enzymes has been shown to improve pentose phosphate pathway fluxes and availability of ribulose-5-phosphate ^30–32^. Despite *TAL1* over-expression in both strains enabling a slight growth increase when spotted onto solid minimal medium with 1, 2 or 4 % methanol (Supplementary Fig. 1), no growth increase was observed when tested in liquid yeast extract methanol medium.

### Liquid growth and ^13^C-methanol tracer analysis confirms methanol assimilation through central carbon metabolism

After observing methanol- and strain-specific growth differences on solid medium (Fig. 1B), methanol-dependent growth of the ‘hybrid’ XuMP-*pTDH3-SFA1*, bacterial RuMP-*pTDH3-SFA1*, and empty vector control strains was tested in liquid medium. No growth was observed with 2 % methanol as the sole carbon source. Previous studies have highlighted the importance of yeast extract for growth on liquid methanol media ^33^. We therefore tested growth with and without 2 % methanol in liquid YNB medium with 0.1 % yeast extract (Fig. 2A). The presence of methanol in the medium resulted in significant (p < 0.05) final OD_600_ increases relative to yeast-extract-only medium of 58 %, 47 %, and 39 % for the ‘hybrid’ XuMP-*pTDH3-SFA1*, bacterial RuMP-*pTDH3-SFA1*, and control strains, respectively. This result identified a methanol-specific growth increase in liquid medium containing yeast extract. In contrast to the tests on solid methanol YNB medium (Fig. 1B), comparing the growth of the different strains on liquid yeast extract methanol medium revealed only marginal and non-significant differences. The presence of yeast extract appeared to largely eliminate the strain-specific differences that were observed on solid YNB-methanol plates, which is consistent with the trend observed on solid YNB-methanol plates with 0.1% yeast extract (Fig. 1B).

**Fig. 2.**
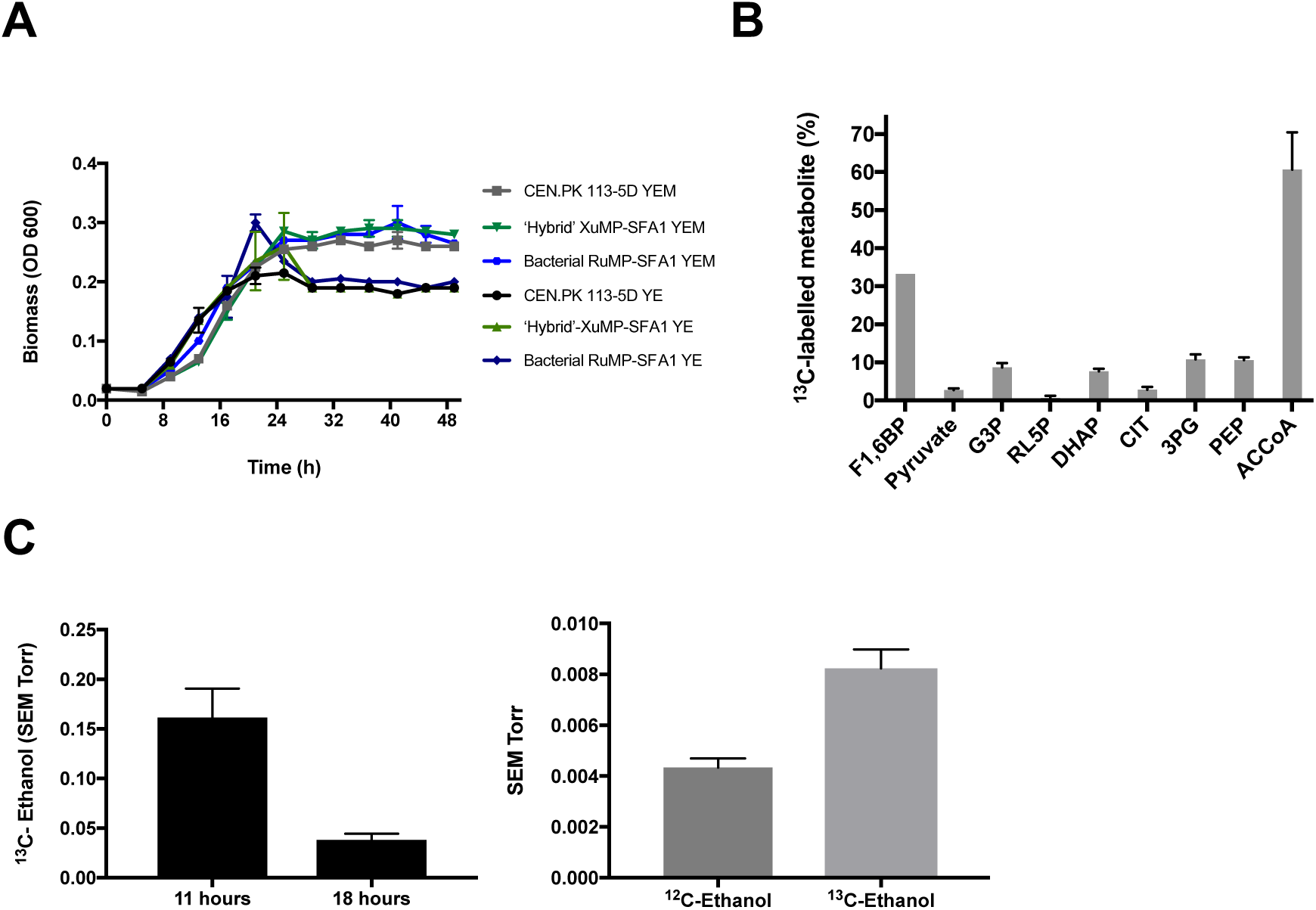
^13^C-methanol fermentations confirm methanol assimilation in *S. cerevisiae*. **A**. Growth profiles of CEN.PK 113-5D (the empty vector control), the ‘hybrid’ XuMP*-pTDH3-SFA1* and the bacterial RuMP-*pTDH3-SFA1* strain grown in liquid YNB medium with 0.1 % yeast extract (YE) or with 0.1 % yeast extract and 2 % methanol (YEM). Data points represent the average of two biological replicates and error bars are the standard deviation. **B.** Percentage of ^13^C-labelled intracellular metabolites in CEN.PK 113-5D, metabolites are universally (fully) labelled with ^13^C. F1,6BP, fructose 1,6-bisphosphate; G3P, glyceraldehyde-3-phosphate; RL5P, ribulose-5-phosphate; DHAP, dihydroxyacetone phosphate; CIT, citrate; 3PG, 3-phosphoglyceric acid; PEP, phosphoenolpyruvate; ACCoA, acetyl-coenzyme A. Data points represent the average of two biological replicates and error bars are the standard deviation. **C.** ^13^C-ethanol was produced by CEN.PK 113-5D. The signal intensity was normalised to the inert gas nitrogen, and then to biomass. Data shows the average ^13^C-ethanol intensity at 47 amu for two biological replicates during independent scanning cycles using a Hiden HPR-20-QIC mass spectrometer. Error bars are the standard deviation of the ^13^C-ethanol intensity. Ratio of ^12^C-ethanol and ^13^C-ethanol produced by CEN.PK 113-5D at 18 hours. The signal intensity was normalised to the inert gas nitrogen, and then to biomass for each strain. Data shows the average ^12^C-ethanol and ^13^C-ethanol intensity at 31 and 33 amu, respectively for two biological replicates during independent scanning cycles using a Hiden HPR-20-QIC mass spectrometer. Error bars are the standard deviation of the ^12^C-ethanol or ^13^C-ethanol intensity. SEM, Secondary Electron Multiplier (SEM); amu, atomic mass unit.

Due to the clear methanol response in all strains, and the marginal differences observed between the wild-type and the engineered strains in liquid medium, the empty vector control strain (CEN.PK 113-5D) was chosen for ^13^C-methanol tracer analysis. Furthermore, as the effect of methanol on the wild-type was unexpected, we first chose to verify methanol utilisation using ^13^C-tracer analysis in this strain. For ^13^C-methanol tracer analysis, bioreactor off-gases such as ^13^C-methanol, ^13^C-CO_2_, CO_2_, ^13^C-ethanol, and ethanol were measured in real-time using a mass spectrometer connected to the bioreactors. Intracellular metabolites were extracted and analysed using Liquid Chromatography-Mass Spectrometry (LC-MS). When the ratio of labelled to unlabelled CO_2_ was compared at the maximum ^13^C-CO_2_ production point, 80 % of total CO_2_ was produced as ^13^C-CO_2_ (Supplementary Fig. 2A). Surprisingly, CEN.PK 113-5D also produced ^13^C-ethanol (Fig. 2C), and ^13^C-methanol tracer analysis furthered confirmed assimilation with universally ^13^C-labelled intracellular metabolites involved in the pentose phosphate pathway, glycolysis, and the TCA cycle (Fig. 2B). Methanol assimilation through central carbon metabolism was therefore identified in the wild-type strain with the detection of various intracellular metabolites including ^13^C-fructose 1,6-bisphosphate and ^13^C-pyruvate, which could only arise through the conversion of ^13^C-methanol through central carbon metabolism (Fig. 2B/C). This result showed for the first time that *S. cerevisiae* has a previously undiscovered native capacity for methanol assimilation. The same ^13^C-methanol tracer analysis was performed for the bacterial RuMP-*pTDH3-SFA1* strain, which revealed a similar metabolic profile as the one observed for CEN.PK 113-5D (Supplementary Fig. 2).

### Adaptive laboratory evolution of native methanol assimilation in CEN.PK 113-5D

As methanol assimilation in *S. cerevisiae* was previously undiscovered, Adaptive Laboratory Evolution (ALE) was applied to characterise and optimise this metabolic response in the wild-type CEN.PK 113-5D strain. The wild-type strain was selected for optimisation as only slight differences in the metabolic profiles of the wild-type strain compared to the bacterial RuMP-*pTDH3-SFA1* strain were observed. The ALE strategy consisted of three independent lineages of CEN.PK 113-5D with an empty pRS416 vector grown with 1x YNB medium with either yeast extract or yeast extract plus 2 % methanol. Yeast-extract-only medium was included to track any improvements in fitness specific to yeast extract. Cultures were grown in baffled shake flasks with alternating passages on 1 % glucose or yeast extract with or without methanol (Fig. 3A). All six independent lineages were grown for 230 generations until a biomass (OD_600_) increase in yeast extract methanol medium was observed. The six evolved strains were re-sequenced to identify putative mutations leading to the phenotype. No relevant mutations were observed in the three lineages evolved in yeast-extract-only medium. In contrast, the three lineages evolved in yeast extract methanol medium all had point mutations at different positions in *YGR067C*, an uncharacterised transcription factor with a DNA-binding domain similar to that of *ADR1* (alcohol dehydrogenase II synthesis regulator). *ADRI* encodes a transcription factor which is involved in the expression of glucose-repressed genes, ethanol metabolism, and peroxisomal proliferation ^34, 35^. All three independent mutations resulted in premature stop codons, which would result in truncation of the *YGR067C* protein (Fig. 3B). To test if truncation of the *YGR067C* transcription factor was responsible for improved growth on methanol, CRISPR-Cas9-mediated homologous recombination was used to introduce a stop-codon in *YGR067C* of the wild-type strain as observed in the evolved lineage C (Fig. 3B). This reconstructed CEN.PK 113-5D strain was referred to as reconstructed EC. The three evolved lineages, the parental CEN.PK 113-5D, and reconstructed EC strains were grown on yeast extract methanol medium to analyse growth differences (Fig. 3C). The evolved lineage A strain had a final biomass increase of 37 % in the presence of methanol compared to the parental CEN.PK 113-5D strain while the evolved lineage B and C strains have a 22 and 44 % increase respectively. Importantly, the reconstructed EC strain has the same growth profile and final biomass increase (44 %) as the evolved lineage C strain compared to the parental strain, indicating that the truncated transcription factor is responsible for the beneficial response to methanol.

**Fig. 3.**
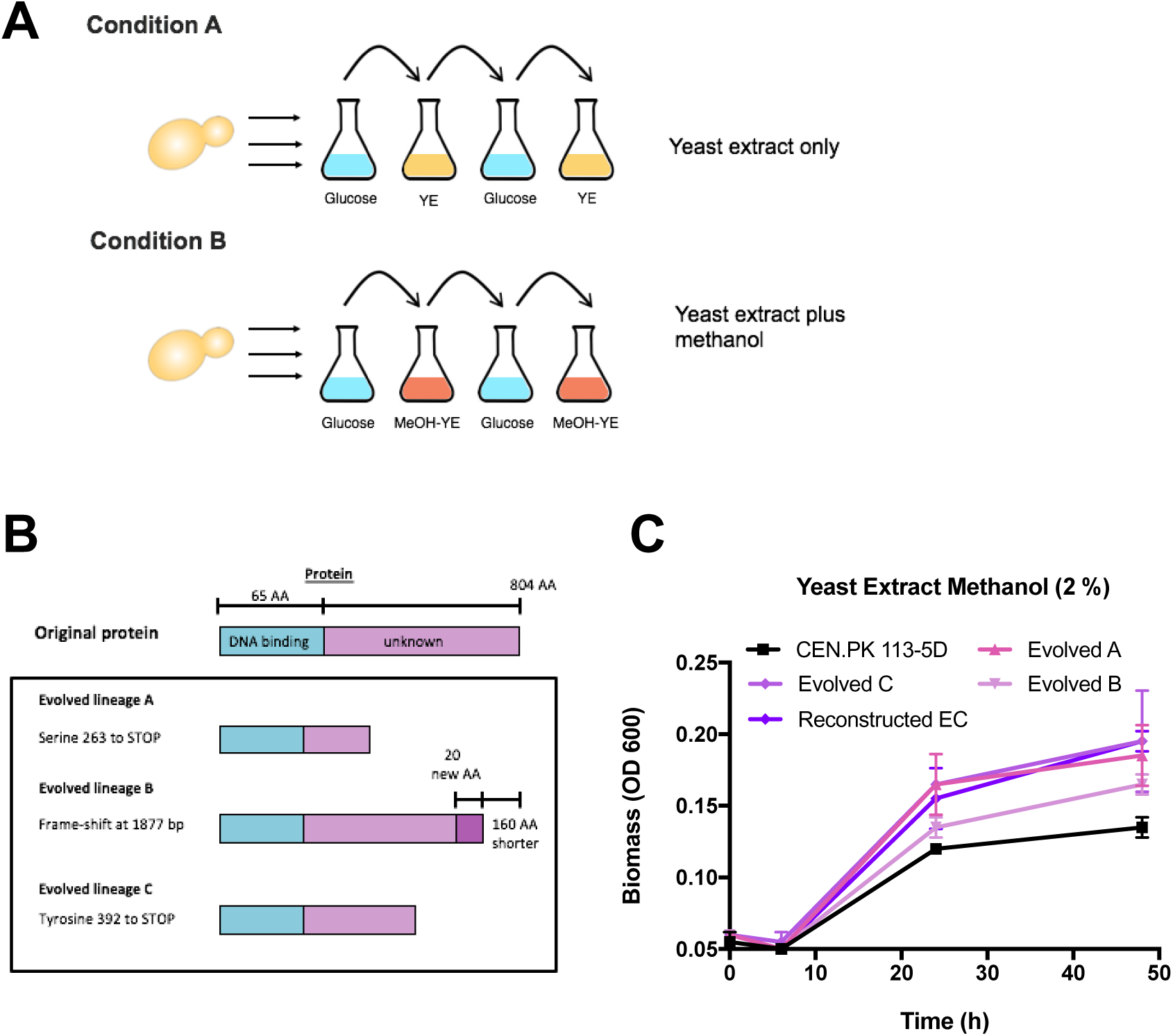
Adaptive Laboratory Evolution to improve native methanol assimilation in CEN.PK 113-5D. **A.** Schematic of the ALE design. Three independent lineages of CEN.PK 113-5D were grown in baffled shake flasks in condition A or B for 230 generations. Under condition A, cultures were passaged from YNB medium without amino acids and 1 % glucose (24 hours) to YNB medium without amino acids and 0.1 % yeast extract (48 hours). Under condition B, cultures were passaged from YNB medium without amino acids and 1 % glucose to YNB medium without amino acids, 2 % methanol and 0.1 % yeast extract (48 hours). **B.** Schematic of the mutations in *YGR067C* from the three evolved lineages grown under condition B, and the changes they caused to the protein, all three mutations theoretically led to truncations. **C.** Growth profiles of CEN.PK 113-5D, the three evolved lineages in condition B, and the reconstructed CEN.PK 113-5D strain with the mutation observed in the evolved lineage C. Strains were grown in liquid YNB medium with 0.1 % yeast extract and 2 % methanol. Data points represent the average of three biological replicates and error bars are the standard deviation.

### Reconstructed EC strain characterisation and ^13^C-methanol tracer analysis

To characterise the effect the reconstructed EC strain had on native methanol metabolism in *S. cerevisiae*, CEN.PK 113-5D and the reconstructed EC strain were grown in bioreactors with 2 % ^13^C-methanol and 0.1 % yeast extract. As previously noted (Fig. 3C), a growth advantage was observed in the reconstructed EC strain (Fig. 4A), which reached a higher final biomass compared to CEN.PK 113-5D. Both strains produced 80 % of total CO_2_ as ^13^C-CO_2_ (Fig. 4B). However, the reconstructed EC strain produced significantly less ^13^C-ethanol, particularly at 11 hours (Fig. 4C). We hypothesized that the reconstructed EC strain could be redirecting methanol into biomass constituents and thus reducing ethanol production. ^13^C-methanol tracer analysis revealed striking differences when intracellular metabolites were compared. The reconstructed EC strain had a higher percentage of universally ^13^C-labelled intracellular metabolites compared to CEN.PK 113-5D (Fig. 5). A ∼3-fold increase in ^13^C-labelled glyceraldehyde-3-phosphate and dihydroxyacetone phosphate was observed as well as increased ^13^C-labelling of metabolites involved in lower glycolysis, including 3-phosphoglyceric acid, phosphoenolpyruvate, and pyruvate. ^13^C-labelled pyruvate increased from 2.7 % in CEN.PK113-5D to 15.4 % in the reconstructed EC strain. Higher ^13^C-labelling was also observed for metabolites in the pentose phosphate pathway but the proportion of ^13^C-labelled metabolites was still low in both strains (less than 5 %). An interesting observation was that CEN.PK 113-5D had a higher proportion of ^13^C-labelled acetyl-CoA compared to the reconstructed EC strain, 67.6 % compared to 46.6 %. Finally, no unlabelled or ^13^C-labelled α-ketoglutarate, fumarate, malate, oxaloacetate or glyoxylate were detected.

**Fig. 4.**
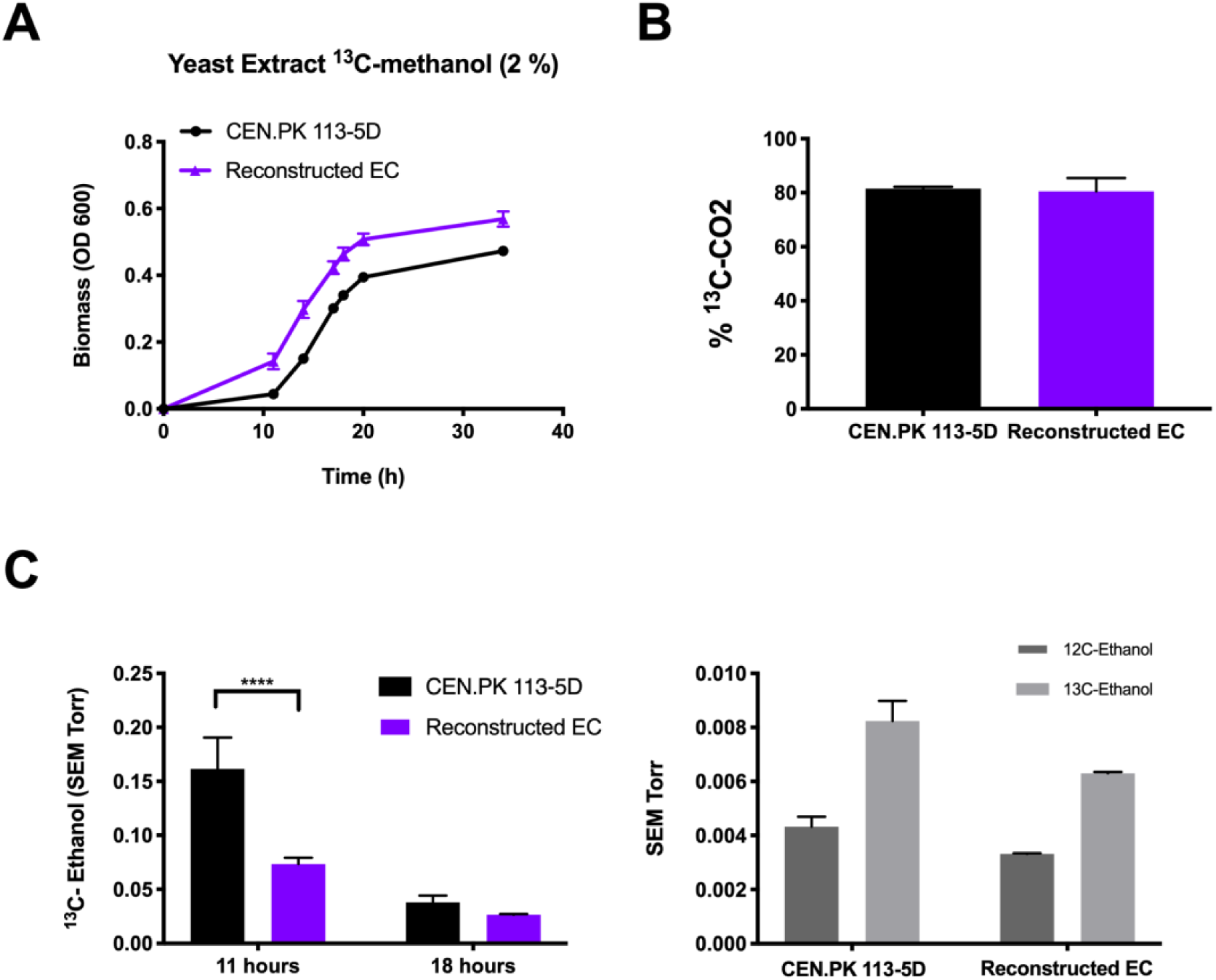
^13^C-methanol fermentations to characterise the reconstructed evolved strain. A. Growth profile of CEN.PK 113-5D and the reconstructed evolved strain grown in liquid YNB medium with 2 % ^13^C-methanol supplemented with 0.1 % yeast extract cultures in bioreactors. **B.** Percentage of ^13^C-CO_2_/CO_2_ production in yeast extract ^13^C-methanol (2 %) medium **C.** ^13^C-ethanol was produced by CEN.PK 113-5D and by the reconstructed evolved strain. The signal intensity was normalised to the inert gas nitrogen, and then to biomass for each strain. Data shows the average ^13^C-ethanol intensity at 47 amu for two biological replicates during independent scanning cycles using a Hiden HPR-20-QIC mass spectrometer. Error bars are the standard deviation of the ^13^C-ethanol intensity. Ratio of ^12^C-ethanol and ^13^C-ethanol produced at 18 hours. The signal intensity was normalised to the inert gas nitrogen, and then to biomass for each strain. Data shows the average ^12^C-ethanol and ^13^C-ethanol intensity at 31 and 33 amu, respectively for two biological replicates during independent scanning cycles using a Hiden HPR-20-QIC mass spectrometer. Error bars are the standard deviation of the ^12^C-ethanol or ^13^C-ethanol intensity. SEM, Secondary Electron Multiplier; amu, atomic mass unit. Data points represent the average of two biological replicates and error bars are the standard deviation.

**Fig. 5.**
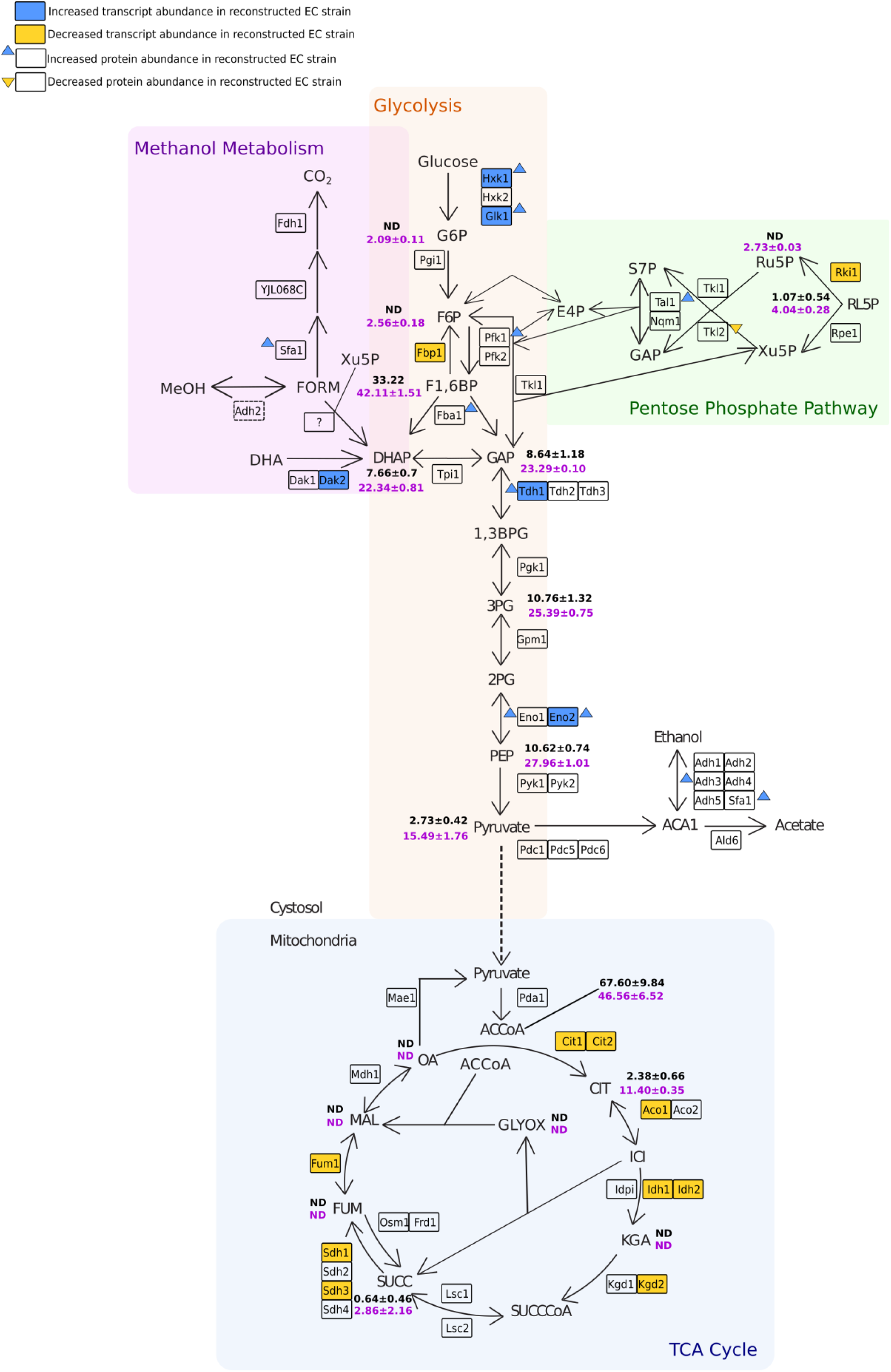
Characterisation of the reconstructed evolved strain at the metabolite, transcriptome and proteome level compared to the parental strain during ^13^C-methanol fermentations. Percentage of ^13^C-labelled intracellular metabolites in CEN.PK 113-5D (black) and the reconstructed evolved strain (purple) at exponential phase. Metabolites are universally (fully) labelled with ^13^C. Data points represent the average of two biological replicates with standard deviation. Transcript and protein abundance in the reconstructed evolved strain compared to the parental strain. Genes with up- or down-regulated fold-changes and adjusted p-values less than 0.01 are coloured blue or yellow respectively. Increased or decreased protein abundance and adjusted p-values less than 0.05 is shown in blue or yellow arrows respectively. RNA and protein samples were taken at stationary phase. ND; not detected. The metabolic map was adapted from ^39^.

To further characterise the changes in the reconstructed EC strain, global transcript and protein levels were compared with the parental strain (Fig. 5). During growth on 2 % ^13^C-methanol and 0.1 % yeast extract, 243 transcripts were found to be significantly differentially expressed in the reconstructed EC strain (adjusted p < 0.01) with 111 genes down-regulated and 132 genes up-regulated (Supplementary File 1). Gene-list analysis of the down-regulated genes showed Gene Ontology (GO) pathway enrichment of the TCA cycle, respiration, and the glyoxylate cycle. No pathway enrichment was found for the up-regulated genes in the reconstructed EC strain but GO process enrichment was found for ‘carbohydrate transmembrane transport’, ‘carbohydrate metabolic process’, ‘glycolytic process’ and ‘pyruvate metabolic process’, among others. The proteomics analysis showed 103 proteins were significantly altered in abundance in the reconstructed EC strain relative to the parent (adjusted p < 0.05; Supplementary File 2) with 84 proteins increased and 19 proteins decreased. Gene-list analysis of proteins with increased expression showed pathway enrichment of the ‘superpathway of glucose fermentation’ while the proteins with decreased abundance showed GO process enrichment only for ‘trehalose metabolic process’. When comparing the proteome of the reconstructed EC strain to that of the parental CEN.PK 113-5D strain, 79 proteins were not identified in the proteome of the parental strain, while 14 proteins were not identified in the reconstructed EC strain (Supplementary File 2).

Both transcriptomics and proteomics indicated there were significant changes in central carbon metabolic pathway expression between the evolved and parental strain. Five genes involved in glycolysis had higher transcript abundance in the reconstructed EC strain compared to CEN.PK 113-5D (*HXK1*, *GLK1*, *DAK2*, *TDH1* and *ENO2*) while *FBP1*, coding for fructose-1,6-bisphosphatase, a key enzyme involved in gluconeogenesis ^36, 37^, had lower transcript abundance. Hxk1p, Glk1p, Tdh1p, and Eno2p also had increased protein abundance, as well as Pfk1p, suggesting glycolysis rather than gluconeogenesis is predominantly occurring during growth on methanol in the reconstructed EC strain. The increased transcript abundance of the 6-phosphofructo-2-kinase glycolysis regulator *PFK27* ^38^ in the reconstructed EC strain also supports this concept. Similar to the metabolite profile of the reconstructed EC strain, lower transcript abundance was observed for nine genes involved in the TCA cycle (*CIT1, CIT2*, *ACO1*, *IDH1*, *IDH2*, *KGD2*, *SDH1*, *SDH3* and *FUM3*; Fig. 5), suggesting the TCA cycle is downregulated during growth on methanol. Sfa1p, which is involved in native formaldehyde detoxification and alcohol oxidation also had increased protein abundance in the reconstructed EC strain. This is consistent with the growth benefit we observed from *SFA1* over-expression (Fig. 1B & Fig. 2A). Finally, the pentose phosphate pathway showed interesting results, with *RKI1* having lower transcript abundance and Tkl2p lower protein abundance while Tal1p and Fba1p had higher protein abundance. This could suggest transaldolase instead of transketolase is the preferred enzyme to yield ribose-5-phosphates through the non-oxidative branch of the pentose phosphate pathway in the reconstructed EC strain.

After characterising the reconstructed EC strain and identifying an improvement in native methanol assimilation, the synthetic pathways highlighted in Fig. 1 were engineered into the reconstructed EC strain (Supplementary Fig. 3). The synthetic pathways improved growth in the reconstructed EC strain as they did in the parental CEN.PK 113-5D strain (Fig. 1B & 2A). The ‘hybrid’ XuMP and bacterial RuMP with over-expression of *SFA1* had increased growth on solid minimal media with increasing concentrations of methanol compared to the reconstructed EC strain (Supplementary Fig. 3A), however, no growth advantage was observed when the reconstructed EC strain containing the bacterial RuMP-*pTDH3-SFA1* pathway was grown in bioreactors with yeast extract and ^13^C-methanol (2 %) medium (Supplementary Fig. 3B). The presence of yeast extract results in slight, non-significant strain-specific differences, as was previously observed when comparing growth on solid methanol media with liquid yeast extract methanol media (Fig. 1B & Fig. 2A). Finally, both the reconstructed EC strain and the reconstructed EC strain containing the bacterial RuMP-*pTDH3-SFA1* pathway show a similar metabolic profile (Supplementary Fig. 3C/D/E) suggesting no meaningful differences are apparent between the strains when grown in liquid medium, although significant methanol-specific differences were apparent on solid medium.

### *ADH2* and *ACS1* deletion reduces methanol-specific growth

Eight genes were selected to analyse their potential role in native methanol assimilation in the reconstructed EC strain. Firstly, the gene coding for an alcohol dehydrogenase 2 (*ADH2*) was selected as in a separate study it was significantly up-regulated in response to methanol ^40^ and due to the promiscuity of alcohol dehydrogenases in *S. cerevisiae*, it could be oxidising methanol to formaldehyde. *CAT8*, coding for a transcription factor involved in de-repressing genes during growth on non-fermentable carbon sources and thought to regulate the mutated transcription factor Ygr067cp ^41^ (Fig. 3B) was chosen to analyse its effect on methanol growth. The serine hydroxymethyltransferase gene (*SHM1*) was chosen for deletion as another possibility for C1 carbon assimilation in *S. cerevisiae* involves formaldehyde detoxification to formate and then assimilation through the glycine cleavage complex ^42^. Shm1p is responsible for converting serine to glycine and 5,10-methylenetetrahydrofolate and would be required if biomass formation was due to formaldehyde detoxification and subsequent formate assimilation. The second set of genes selected for analysis were *DAK2*, *ENO1*, *ENO2, WSC3*, and *ACS1*. *DAK2* and *ENO2*, coding for a dihydroxyacetone kinase and a phosphopyruvate hydratase, respectively, were selected as they had higher transcript abundance in the reconstructed EC strain compared to the parent and would be required for important steps during glycolysis (Fig. 5). *ENO1* was also selected as it is a paralog of *ENO2* ^43^. *WSC3* was selected as a putative methanol sensor since a study by Ohsawa et al. (2017) identified Wsc1p and Wsc3p from *P. pastoris* as sensors involved in its native methanol response, with Wsc3p from *P. pastoris* having 67.4 % amino acid sequence similarity with Wsc3p from *S. cerevisiae* ^44^. Wsc3p in *P. pastoris* is thought to act as a methanol sensor involved in activating a signalling cascade for the expression of methanol-inducible genes. Finally, *ACS1* coding for an Acetyl-CoA synthetase was selected as a study by Oshawa et al. (2018) analysed the role of *ACS1* in *P. pastoris* and found that its involved in ethanol repression, which is needed for the expression of methanol-induced genes ^45^.

Single deletion strains of the aforementioned genes, except for *DAK2*, were constructed in the reconstructed EC strain and tested by spotting the strains onto solid minimal YNB medium with 2 % glucose, no additional carbon source, or increasing methanol concentrations (Fig. 6 & Supplementary Fig. 4). The reconstructed EC strain *Δadh2* grew similarly to the reconstructed EC strain in minimal medium with 2 % glucose or no additional carbon source (1 x YNB) but growth was dramatically reduced on media containing methanol (1 to 3 %), suggesting it is needed for methanol oxidation to formaldehyde. When *Δcat8* was deleted from the reconstructed EC strain, growth was only observed with 2 % glucose as the carbon source, and its deletion dramatically decreased growth on methanol and 1 x YNB, as previously reported ^46^. Any potential role of *CAT8* in methanol assimilation could therefore not be analysed using this method. Deletion of *SHM1* had no effect on growth compared to the reconstructed EC strain on media with either glucose or methanol, suggesting it is not involved in the native methanol assimilation pathway. Finally, the reconstructed EC strains with *Δeno2, Δeno1, Δacs1, Δwsc3* showed inconclusive results (Supplementary Fig. 4). The reconstructed EC strain with *Δeno2* and *Δwsc3* had slight growth reductions on media with methanol but not glucose, *Δacs1* showed almost no growth when methanol was present at higher concentrations (2 and 3 %), and *Δeno1* showed no growth differences between media.

**Fig. 6.**
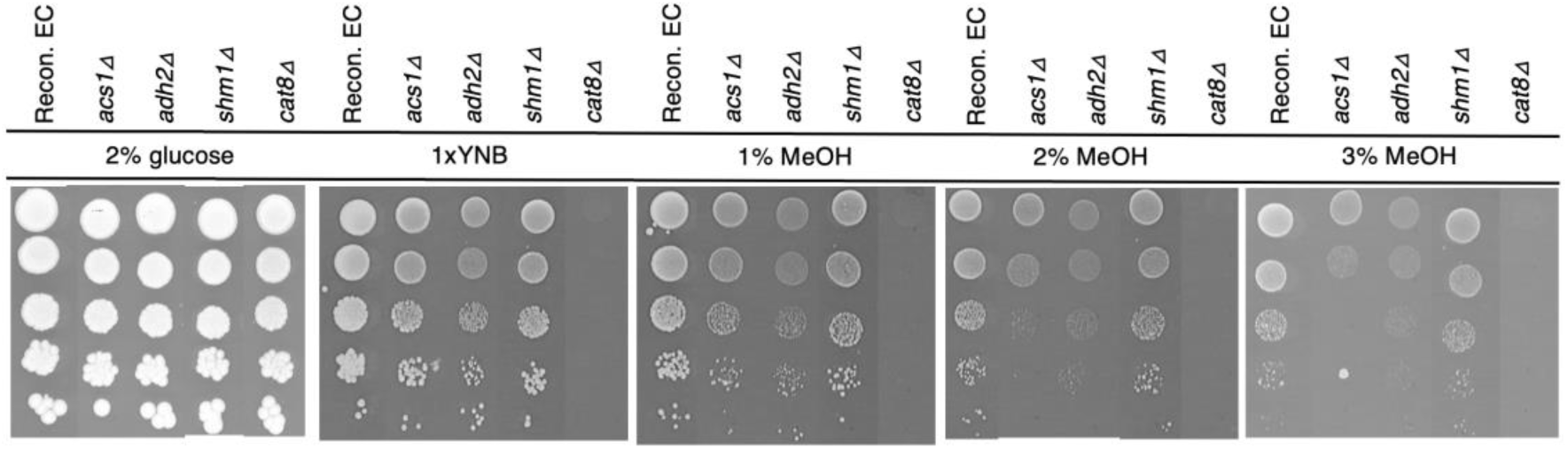
Growth in methanol of different gene deletions to test their putative involvement *in S. cerevisiae*’s native methanol assimilation. Growth on solid 1x Yeast Nitrogen Base medium with different carbon sources was tested using serial 10-fold dilutions of the reconstructed evolved strain with an empty vector or *ACS1, ADH2*, *SHM1* or *CAT8* deletions. Yeast Nitrogen Base (YNB), Yeast Extract (YE), Methanol (MeOH). Images were taken after incubating at 30 °C for 6 days.

## Discussion

As methanol is emerging as an important C1 feedstock, we engineered the XuMP and RuMP methanol assimilation pathways in *S. cerevisiae* (Fig. 1A), and analysed growth on methanol as well as ^13^C-metabolite production from ^13^C-methanol. The XuMP pathway was targeted either to the peroxisome or to the cytosol, where a ‘hybrid’ version of the pathway was designed. A subtle growth improvement only on solid 1 % methanol medium was observed when the enzymes were targeted to the peroxisome by a PTS1 signal (Fig. 1B). This pathway is based on *P. pastoris* metabolism, in which peroxisome proliferation plays a major role during growth on methanol, a common characteristic of other methylotrophic yeasts ^29, 47^. However, peroxisome proliferation in *S. cerevisiae* is limited to specific stress responses such as fatty acid oxidation and requires the regulation of multiple genes ^48^. It is possible that peroxisome proliferation was a limiting step for the function of the XuMP pathway as peroxisome proliferation in response to methanol has not evolved in *S. cerevisiae*. In conclusion, a XuMP methanol assimilation pathway targeted to the peroxisome did not appear to be a viable option for synthetic methylotrophy in *S. cerevisiae*. This contradicts a previous report by Dai et al. where up to 12 % improvement on growth was recorded in yeast extract methanol (1 %) medium after introduction of the yeast XuMP pathway (Aox1p-Das1p-Cat1p-Dak1p or Aox1p-Das2p-Cat1p-Dak1p) ^49^.

The second, cytosol localised, ‘hybrid’ XuMP pathway utilised a bacterial NAD^+^-dependent Mdh alongside the *P. pastoris DAS1*. The third ‘bacterial RuMP’ pathway used enzymes from methylotrophic bacteria (Mdh and Hps-Phi). An NAD^+^-dependent methanol dehydrogenase was chosen to oxidise methanol to formaldehyde (first step on the pathway), as it has been identified as having higher affinity for methanol and the most efficient conversion compared to other methanol dehydrogenases, especially the variant from *B. stearothermophilus* ^16, 18^. To assimilate formaldehyde to biomass, enzymes utilising two different cyclic pathways were chosen, Das1p (XuMP pathway) or Hps and Phi (RuMP pathway). These two alternatives were tested as they use different co-substrates for formaldehyde assimilation (xylulose-5-phosphate or ribulose-5-phosphate) and produce either glyceraldehyde-3-phosphate or fructose-6-phosphate as the precursors for biomass formation (Fig. 1A). Both pathways enabled a greater capacity for growth on solid minimal methanol medium relative to an empty vector control strain, with the bacterial RuMP pathway being superior to the ‘hybrid’-XuMP (Fig. 1B). This result is supported by an *in silico* flux balance analysis performed by Comer et al., which analysed different C1 carbon sources and also found the RuMP pathway to be the more promising option in *S. cerevisiae* ^50^. Moreover, a recent study in *Yarrowia lipolytica* containing an *FLD* (formaldehyde dehydrogenase) deletion found that formaldehyde tolerance was recovered when *B. methanolicus* Hps was introduced, while *P. pastoris* Das1p (with or without peroxisomal targeting) was only capable of partially restoring formaldehyde tolerance ^51^.

An interesting development while testing the synthetic methylotrophic pathways in yeast was the identification of formaldehyde detoxification as an important component of methanol-dependent growth improvement. Studies on methylotrophic species have previously highlighted the importance of the balance between assimilation and dissimilation of formaldehyde. Over-expression of *SFA1*, the first gene in the native formaldehyde detoxification pathway, was therefore tested alongside the three synthetic pathways. Growth on methanol was significantly improved in all strains upon introduction of an over-expressed *SFA1* (Fig. 1B). Likewise, methanol assimilation in solid minimal YNB-methanol media was identified in both the ‘hybrid’ XuMP and bacterial RuMP strains by comparing growth against an empty vector control strain on increasing and toxic concentrations of methanol. Recent work has also highlighted the importance of formaldehyde dissimilation through the engineering of synthetic methylotrophy in *E. coli*. For example, Müller et al. detected an overall higher percentage of total ^13^C-labelling when they assessed methanol assimilation in a wild-type background compared to a genetic background unable to dissimilate formaldehyde (Δ*frmA*; 39.4 % versus 25.2 %), results that also contradicted their *in silico* simulations where formaldehyde detoxification was not found to be essential for the conversion of methanol ^16^. Woolston et al. also investigated the improvement of formaldehyde assimilation in a synthetic methylotrophic *E. coli* strain. They found that the unfavourable thermodynamics of the Mdh enzyme need to be overcome to maintain the forward reaction direction of methanol oxidation ^21^. One way to achieve this is to maintain low intracellular formaldehyde levels through formaldehyde dissimilation by over-expressing formaldehyde dehydrogenase. This observation could explain why Müller et al. saw higher ^13^C-labelling levels in a wild-type background that was able to dissimilate formaldehyde, and together these observations support the importance of formaldehyde dissimilation for methanol assimilation in *S. cerevisiae*.

Methanol-specific growth of the control, ‘hybrid’ XuMP-*pTDH3-SFA1* and bacterial RuMP-*pTDH3-SFA1* strains, was observed in liquid yeast extract medium with methanol (Fig. 2A). However, strain-specific differences in liquid yeast extract methanol medium were dramatically diminished relative to those observed on minimal solid methanol medium, as observed when yeast extract was added on solid medium (Fig. 1B). ^13^C-labelled methanol was therefore used to analyse the methanol response of CEN.PK 113-5D with an empty vector control in liquid medium. Fermentations with 2 % ^13^C-methanol were performed and quite surprisingly, ^13^C-ethanol and ^13^C-labelled intracellular metabolites were produced (Fig. 2B/C), demonstrating methanol assimilation through central carbon metabolism. Importantly, *S. cerevisiae* has always been recognised as a non-native methylotroph and no previous growth on- or assimilation of-methanol has been recorded in the literature. The identification of fully ^13^C-labelled metabolites from ^13^C-methanol here clearly shows for the first time that methanol can be assimilated through central carbon metabolism in *S. cerevisiae*.

After identifying native methanol assimilation in the wild-type, CEN.PK 113-5D strain, we decided to focus on characterising and optimising this pathway. To optimise methanol assimilation, an ALE strategy was designed where independent lineages of CEN.PK 113-5D were passaged through conditional media (Fig. 3A). An evolutionary approach was chosen as methanol assimilation in *S. cerevisiae* utilises an uncharacterised pathway and evolution can be used to derive non-obvious solutions to complex and partially characterised systems. Furthermore, laboratory evolution experiments have been successful in optimising consumption of C1 carbon sources such as methanol and CO_2_ in *E. coli*, and CO_2_ in *P. pastoris* ^15, 52, 53^. After 230 generations, growth was increased in liquid yeast extract methanol medium with whole-genome sequencing revealing mutations in an uncharacterised putative transcription factor *YGR067C* that lead to truncations of the protein (Fig. 3B). The mutation observed in the evolved lineage C was reverse engineered into CEN.PK 113-5D, and the phenotype was restored (Fig. 3C), confirming *YGR067C* is responsible for the optimised growth on methanol. Little is known about Ygr067cp, it shares an identical DNA-binding domain with Adr1p and is regulated by Cat8p ^35, 41^. Both Adr1p and Cat8p are involved in de-repressing genes needed for growth on non-fermentable carbon sources ^41^, and Adr1p also activates genes involved in peroxisomal biogenesis and proliferation ^35^. Genes under the control of *ADR1* and relative to methanol metabolism include formate dehydrogenases (*FDH1*/*FDH2*), *ADH2*, catalase (*CTA1*) and *PEX11*, a peroxisomal membrane protein ^35^. The truncation of Ygr067cp could have caused the constant expression of genes that are normally repressed, leading to more favourable regulatory conditions and metabolic fluxes for growth on methanol.

To characterise the improved growth on methanol, the reconstructed EC strain was grown in bioreactors with 2 % ^13^C-methanol and compared to the parental strain CEN.PK 113-5D (Fig. 4). The results confirmed the reconstructed EC strain had an improved capacity for methanol assimilation, with higher proportions of ^13^C-labelled intracellular metabolites observed (Fig. 5). Moreover, integrated metabolomics, transcriptomics, and proteomics analyses showed that the reconstructed EC strain has a different metabolic profile compared to the parent, with the TCA cycle being downregulated as well as *FBP1*, an important regulator of gluconeogenesis (Fig. 5). Together with the upregulated genes (*DAK2, TDH1* and *ENO2*) and higher protein abundance of Pfk1p, Fba1p, Tdh1p, and Eno2p, it is likely that a net glycolytic flux is occurring during methanol assimilation in the reconstructed EC strain.

Our results show that truncation of *YGR067C* causes decreased expression of gluconeogenesis, TCA cycle, and glyoxylate cycle genes, which rearranges metabolic fluxes to favour methanol assimilation. The down-regulated transcripts and proteins that we observed in the TCA and glyoxylate cycles fully overlap with those that are normally de-repressed during growth on non-fermentable carbon sources in a *CAT8* and *SNF1* dependent manner ^35, 41^. Given that *CAT8* is known to regulate *YGR067C* ^41^, it is likely that truncation of the *YGR067C* protein facilitates a decoupling of methanol assimilation from the traditional non-fermentable carbon source utilisation phenotype in yeast. At present, it is unclear how these metabolic rearrangements favour methylotrophy in *S. cerevisiae*. One possibility is that glycolytic rather than gluconeogenic fluxes favour the pentose phosphate pathway fluxes necessary for methanol assimilation in the presence of yeast extract. Another point worth noting is that many obligate methylotrophs operate an incomplete, down-regulated TCA cycle through the absence of α-ketoglutarate dehydrogenase activity, which is thought to preclude heterotrophic growth ^54^.

It is unclear how formaldehyde is assimilated in *S. cerevisiae*. However, the high levels of ^13^C-labelling we saw in dihydroxyacetone phosphate, fructose-1-6-bisphosphate, and glyceraldehyde-3-phosphate suggests assimilation occurs via a mechanism similar to the *P. pastoris* XuMP pathway where formaldehyde and xylulose-5-phosphate are converted into dihydroxyacetone and glyceraldehyde-3-phosphate. From the higher transcript abundance of Tal1p and Fba1p, an interesting idea worth exploring is whether the non-oxidative branch of the pentose phosphate pathway is going through different rearrangement reactions than those when *S. cerevisiae* is grown in glucose to recycle the co-substrates needed for formaldehyde assimilation. Transaldolase could be catalysing the rearrangements of fructose-6-phosphate and sedoheptulose-7-phosphate to glyceraldehyde-3-phosphate and erythrose-4-phosphate as it was recently postulated in *P. pastoris* grown on methanol ^25, 55^. To identify specific genes involved in native methanol assimilation, *ADH2* and six other genes were deleted from the reconstructed EC strain. Another study from our group found that *ADH2* was upregulated when *S. cerevisiae* was grown in the presence of methanol ^40^ and when the reconstructed EC strain with an *ADH2* deletion was grown on solid minimal media with increasing concentrations of methanol, growth was severely hindered in all cases, suggesting its role in oxidising methanol to formaldehyde (Fig. 6). We also observed that deletion of acetyl-CoA synthetase (*ACS1*) reduced growth on solid methanol media (Fig. 6). Acs1p is important for growth on non-fermentable carbon sources ^56^, and its homolog is an important regulator of methylotrophic metabolism in *P. pastoris* ^45, 57^. Acs1p is therefore likely to be an important source of acetyl-CoA in *S. cerevisiae* methanol metabolism. The –omics profile of the reconstructed EC strain as well as the identification of Adh2p as the likely first enzyme involved in the native methanol assimilation of CEN.PK 113-5D suggest a *P. pastoris*-like pathway is occurring.

This multi-pathway comparison identified the bacterial RuMP-*pTDH3-SFA1* strain as the best candidate for synthetic methanol assimilation in *S. cerevisiae*. Additionally, native methanol assimilation was identified in *S. cerevisiae* for the first time and optimised through ALE, providing the possibility of exploring the newly discovered native methanol metabolism. Insights into this native methanol response have been elucidated such as methanol oxidation to formaldehyde by Adh2p and altered central carbon metabolism enzyme expression levels. It is possible that *S. cerevisiae* shares some of the characteristics of methylotrophic yeasts like *P. pastoris* and is assimilating methanol through an incomplete version of this methylotrophic pathway (Fig. 1A). Higher transcript abundance of *DAK2* alongside higher protein abundance of Tal1p and Fba1p and high ^13^C-labelling of dihydroxyacetone phosphate also support this hypothesis. Higher biomass formation, a complete understanding of native methylotrophic metabolism, and the elimination of the requirement for yeast extract in liquid methanol medium are the most immediate challenges remaining for the implementation of growth with methanol as the sole carbon source in *S. cerevisiae*. Higher biomass formation is likely limited by the inefficient regeneration of pentose phosphate sugars, which is needed to increase the rate of formaldehyde assimilation. After robust methylotrophy is established in *S. cerevisiae*, a plethora of existing metabolite production pathways could potentially be coupled to a methanol converting ‘platform strain’ using the state-of-the-art genetic tools and the deep physiological characterization available in this model organism. Together, the results from this work represent an exciting step towards using sustainable feedstocks during microbial fermentations for conversion to chemicals, fuels, materials, pharmaceuticals, and foods, and provide the first glimpse of an unexplored metabolic network.

## Methods

### Strains and Plasmids

*S. cerevisiae* plasmids and strains used in this study are shown in Tables 1 and 2, respectively. The genes were codon-optimised for *S. cerevisiae* and synthesised by GenScript USA Inc. DNA manipulation and propagation were performed using standard techniques ^58^. All *S. cerevisiae* transformations were carried out using the lithium acetate method ^59^. *AOX1*, *DAS1*, *mdh*, *hps* and *phi* genes were designed with flanking loxPsym sites to enable compatibility with the synthetic yeast genome SCRaMbLE system, however, this system was not tested during this study ^60^. The synthetic genes were amplified using PCR and cloned into the pRS416 plasmid digested with SmaI using the NEBuilder^®^ HiFi DNA Assembly Master Mix. The *SFA1, TAL1* and *TLK1* genes from *S. cerevisiae* were over-expressed by amplifying the gene and its native terminator from genomic DNA, and cloned under the control of the *TDH3* promoter. These over-expression constructs were cloned into the relevant pRS416 plasmids containing methanol assimilation enzymes, the NEBuilder^®^ HiFi DNA Assembly Master Mix was used for Gibson Assembly ^61^. All Gibson Assemblies were verified via PCR with M13 primers. For multiple site integration of *DAS1*, the gene was cloned into the pBDK1 plasmid between the *PGK1* promoter and terminator flanked by Ty1 delta sites, and transformed into yeast as described previously ^62^, cells were plated on minimal media with 0.5 % glucose and toxic levels of formaldehyde (4 mM) and incubated at 30 °C. To delete *ADH2*, *CAT8*, *SHM1*, *ENO1*, *ENO2*, *WSC3*, *ACS1* and *DAK2* from the reconstructed EC strain, primers amplifying the open reading frame flanked by 500 bp homology regions were designed and the constructs were amplified from the BY4741 knockout collection ^63^. Amplified constructs were then transformed into the reconstructed EC strain and plated on Yeast Extract Peptone Dextrose (YPD) with Geneticin (200 μg / mL; Gibco™ 10131035). Independent colonies were screened via PCR using primers that annealed 200 bp outside of the PCR-generated homology region. Deletion of *DAK2* was never achieved despite multiple attempts. All plasmid maps are available in Supplementary File 3.

**Table 1.** Plasmids used in this study.

| Name | Details | Origin |
| --- | --- | --- |
| **pRS416** | Yeast centromeric plasmid, URA3 marker | Euroscarf <sup>76</sup> |
| **ScMOX3a-416** | *pTDH3-AOX1-CYC1t-pTEF1-DAS1-PHO5t-pSSA1-PYC1-ADH1t*-pRS416 | This study |
| **ScMOX3a-SFA1-416** | *pTDH3-AOX1-CYC1t-pTEF1-DAS1-PHO5t-pSSA1-PYC1-ADH1t-pTDH3-SFA1-SFA1t*-pRS416 | This study |
| **ScMOX3e-416** | *pTEF1-DAS1-PHO5t-pTDH3-mdh-CYC1t*-pRS416 | This study |
| **ScMOX3e2-416** | *pTEF1-DAS1-PHO5t-pTDH3-mdh-CYC1t-pTEF2-DAS2-ADH1t*-pRS416 | This study |
| **ScMOX3e-SFA1-416** | *pTEF1-DAS1-PHO5t-pTDH3-mdh-CYC1t-pTDH3-SFA1-SFA1t*-pRS416 | This study |
| **ScMOX3e-TLK1-416** | *pTEF1-TKL1-TKL1t-pTEF1-DAS1-PHO5t-pTDH3-mdh-CYC1t*-pRS416 | This study |
| **ScMOX3e-TAL1-416** | *pSSA1-TAL1-TAL1t-pTEF1-DAS1-PHO5t-pTDH3-mdh-CYC1t*-pRS416 | This study |
| **ScMOX3g-416** | *pTEF1-hps-ADH1t-pTEF2-phi-PHO5t-pTDH3-mdh-CYC1t*-pRS416 | This study |
| **ScMOX3g-SFA1-416** | *pTEF1-hps-ADH1t-pTEF2-phi-PHO5t-pTDH3-mdh-CYC1t-pTDH3-SFA1-SFA1t*-pRS416 | This study |
| **ScMOX3g-TKL1-416** | *pTEF1-TKL1-TKL1t-pTEF1-hps-ADH1t-pTEF2-phi-PHO5t-pTDH3-mdh-CYC1t*-pRS416 | This study |
| **ScMOX3g-TAL1-416** | *pSSA1-TAL1-TAL1t-pTEF1-hps-ADH1t-pTEF2-phi-PHO5t-pTDH3-mdh-CYC1t*-pRS416 | This study |

**Table 2.**
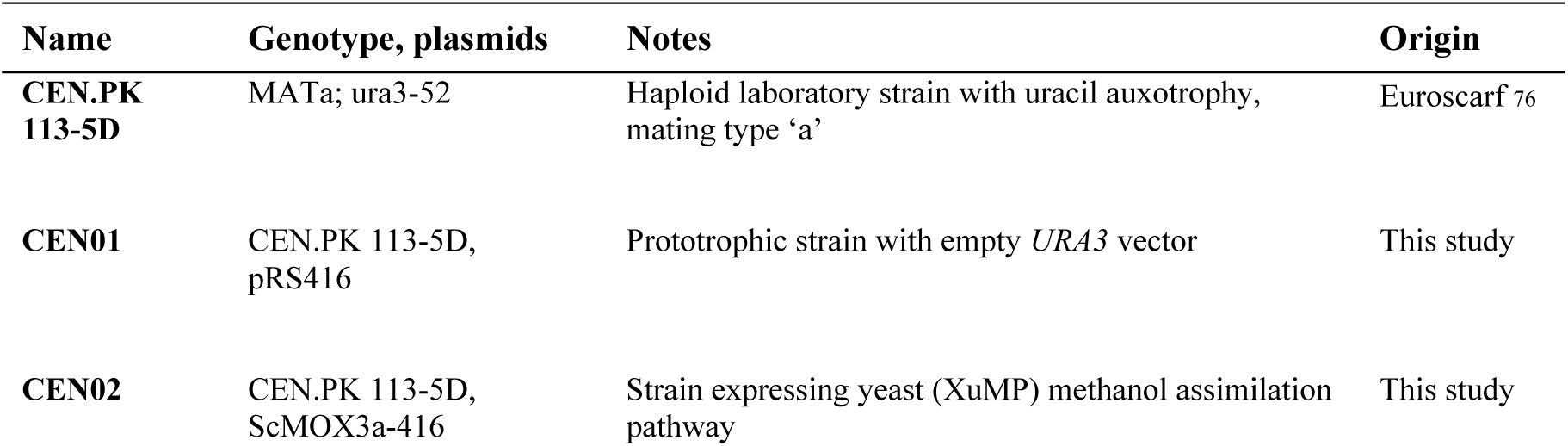
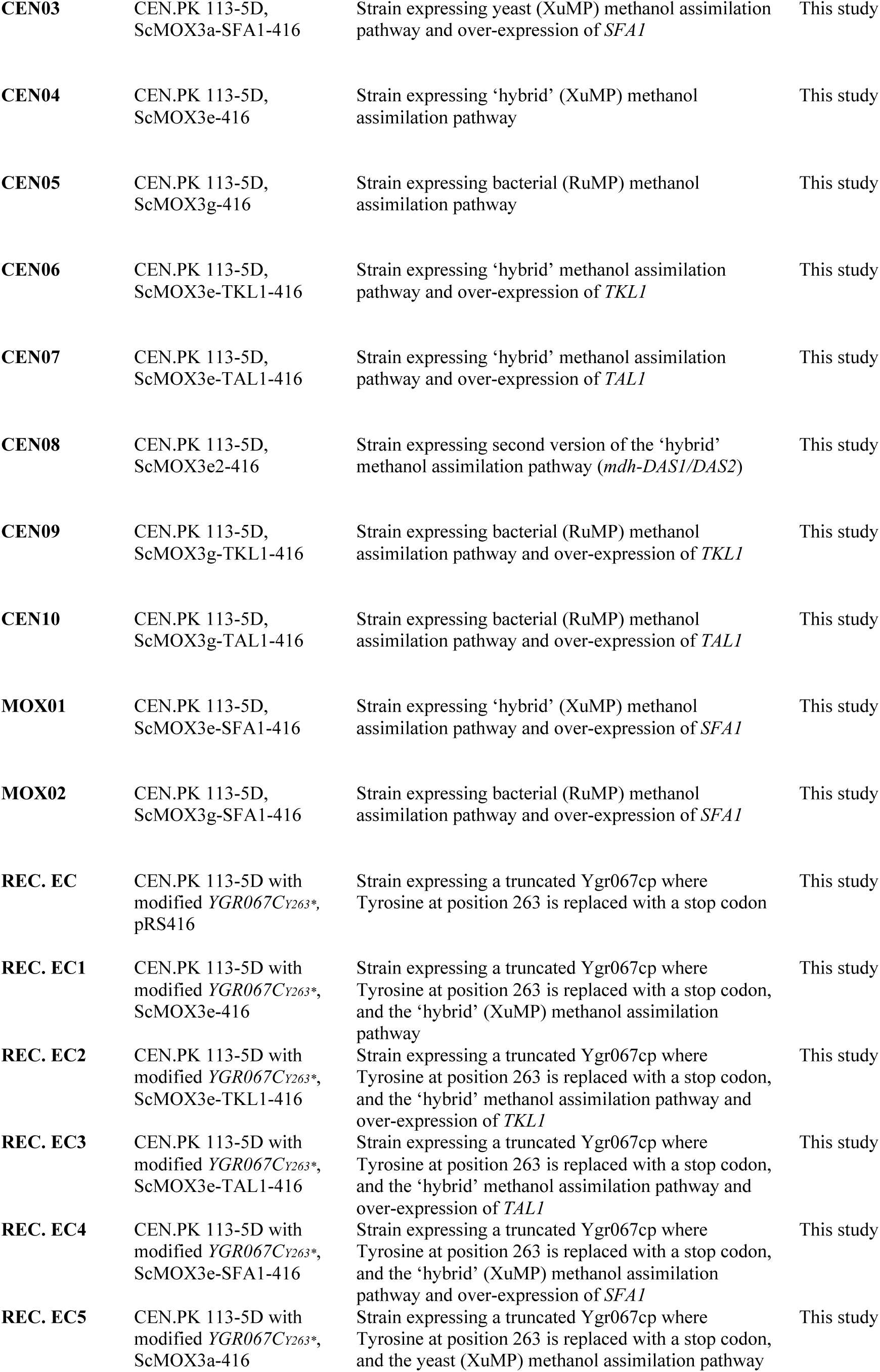
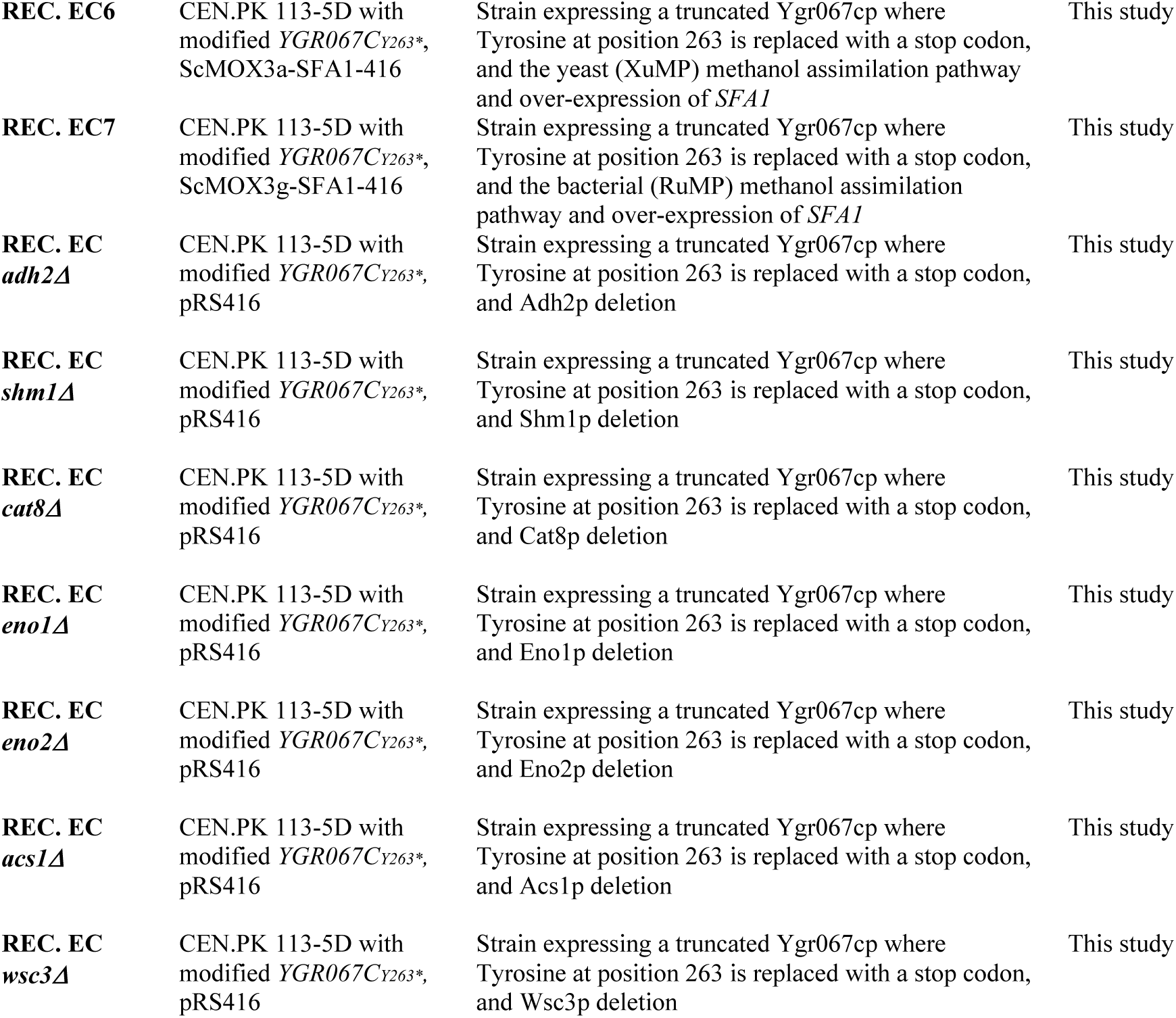
***Saccharomyces cerevisiae* strains used in this study.**

| Name | Genotype, plasmids | Notes | Origin |
| --- | --- | --- | --- |
| **CEN.PK 113-5D** | MATa; ura3-52 | Haploid laboratory strain with uracil auxotrophy, mating type 'a' | Euroscarf <sup>76</sup> |
| **CEN01** | CEN.PK 113-5D, pRS416 | Prototrophic strain with empty *URA3* vector | This study |
| **CEN02** | CEN.PK 113-5D, ScMOX3a-416 | Strain expressing yeast (XuMP) methanol assimilation pathway | This study |
| **CEN03** | CEN.PK 113-5D,<br>ScMOX3a-SFA1-416 | Strain expressing yeast (XuMP) methanol assimilation pathway and over-expression of *SFA1* | This study |
| **CEN04** | CEN.PK 113-5D,<br>ScMOX3e-416 | Strain expressing ‘hybrid’ (XuMP) methanol assimilation pathway | This study |
| **CEN05** | CEN.PK 113-5D,<br>ScMOX3g-416 | Strain expressing bacterial (RuMP) methanol assimilation pathway | This study |
| **CEN06** | CEN.PK 113-5D,<br>ScMOX3e-TKL1-416 | Strain expressing ‘hybrid’ methanol assimilation pathway and over-expression of *TKL1* | This study |
| **CEN07** | CEN.PK 113-5D,<br>ScMOX3e-TAL1-416 | Strain expressing ‘hybrid’ methanol assimilation pathway and over-expression of *TAL1* | This study |
| **CEN08** | CEN.PK 113-5D,<br>ScMOX3e2-416 | Strain expressing second version of the ‘hybrid’ methanol assimilation pathway (*mdh-DAS1/DAS2*) | This study |
| **CEN09** | CEN.PK 113-5D,<br>ScMOX3g-TKL1-416 | Strain expressing bacterial (RuMP) methanol assimilation pathway and over-expression of *TKL1* | This study |
| **CEN10** | CEN.PK 113-5D,<br>ScMOX3g-TAL1-416 | Strain expressing bacterial (RuMP) methanol assimilation pathway and over-expression of *TAL1* | This study |
| **MOX01** | CEN.PK 113-5D,<br>ScMOX3e-SFA1-416 | Strain expressing ‘hybrid’ (XuMP) methanol assimilation pathway and over-expression of *SFA1* | This study |
| **MOX02** | CEN.PK 113-5D,<br>ScMOX3g-SFA1-416 | Strain expressing bacterial (RuMP) methanol assimilation pathway and over-expression of *SFA1* | This study |
| **REC. EC** | CEN.PK 113-5D with<br>modified *YGR067CY263\**,<br>pRS416 | Strain expressing a truncated Ygr067cp where Tyrosine at position 263 is replaced with a stop codon | This study |
| **REC. EC1** | CEN.PK 113-5D with<br>modified *YGR067CY263\**,<br>ScMOX3e-416 | Strain expressing a truncated Ygr067cp where Tyrosine at position 263 is replaced with a stop codon, and the ‘hybrid’ (XuMP) methanol assimilation pathway | This study |
| **REC. EC2** | CEN.PK 113-5D with<br>modified *YGR067CY263\**,<br>ScMOX3e-TKL1-416 | Strain expressing a truncated Ygr067cp where Tyrosine at position 263 is replaced with a stop codon, and the ‘hybrid’ methanol assimilation pathway and over-expression of *TKL1* | This study |
| **REC. EC3** | CEN.PK 113-5D with<br>modified *YGR067CY263\**,<br>ScMOX3e-TAL1-416 | Strain expressing a truncated Ygr067cp where Tyrosine at position 263 is replaced with a stop codon, and the ‘hybrid’ methanol assimilation pathway and over-expression of *TAL1* | This study |
| **REC. EC4** | CEN.PK 113-5D with<br>modified *YGR067CY263\**,<br>ScMOX3e-SFA1-416 | Strain expressing a truncated Ygr067cp where Tyrosine at position 263 is replaced with a stop codon, and the ‘hybrid’ (XuMP) methanol assimilation pathway and over-expression of *SFA1* | This study |
| **REC. EC5** | CEN.PK 113-5D with<br>modified *YGR067CY263\**,<br>ScMOX3a-416 | Strain expressing a truncated Ygr067cp where Tyrosine at position 263 is replaced with a stop codon, and the yeast (XuMP) methanol assimilation pathway | This study |
| **REC. EC6** | CEN.PK 113-5D with modified *YGR067CY263\**, ScMOX3a-SFA1-416 | Strain expressing a truncated Ygr067cp where Tyrosine at position 263 is replaced with a stop codon, and the yeast (XuMP) methanol assimilation pathway and over-expression of *SFA1* | This study |
| **REC. EC7** | CEN.PK 113-5D with modified *YGR067CY263\**, ScMOX3g-SFA1-416 | Strain expressing a truncated Ygr067cp where Tyrosine at position 263 is replaced with a stop codon, and the bacterial (RuMP) methanol assimilation pathway and over-expression of *SFA1* | This study |
| **REC. EC *adh2Δ*** | CEN.PK 113-5D with modified *YGR067CY263\**, pRS416 | Strain expressing a truncated Ygr067cp where Tyrosine at position 263 is replaced with a stop codon, and Adh2p deletion | This study |
| **REC. EC *shm1Δ*** | CEN.PK 113-5D with modified *YGR067CY263\**, pRS416 | Strain expressing a truncated Ygr067cp where Tyrosine at position 263 is replaced with a stop codon, and Shm1p deletion | This study |
| **REC. EC *cat8Δ*** | CEN.PK 113-5D with modified *YGR067CY263\**, pRS416 | Strain expressing a truncated Ygr067cp where Tyrosine at position 263 is replaced with a stop codon, and Cat8p deletion | This study |
| **REC. EC *eno1Δ*** | CEN.PK 113-5D with modified *YGR067CY263\**, pRS416 | Strain expressing a truncated Ygr067cp where Tyrosine at position 263 is replaced with a stop codon, and Eno1p deletion | This study |
| **REC. EC *eno2Δ*** | CEN.PK 113-5D with modified *YGR067CY263\**, pRS416 | Strain expressing a truncated Ygr067cp where Tyrosine at position 263 is replaced with a stop codon, and Eno2p deletion | This study |
| **REC. EC *acs1Δ*** | CEN.PK 113-5D with modified *YGR067CY263\**, pRS416 | Strain expressing a truncated Ygr067cp where Tyrosine at position 263 is replaced with a stop codon, and Acs1p deletion | This study |
| **REC. EC *wsc3Δ*** | CEN.PK 113-5D with modified *YGR067CY263\**, pRS416 | Strain expressing a truncated Ygr067cp where Tyrosine at position 263 is replaced with a stop codon, and Wsc3p deletion | This study |

### Media and growth conditions

*Escherichia coli* DH5α cells were used for plasmid propagation/storage and grown in lysogeny broth media (1 % tryptone, 0.5 % yeast extract, 1 % NaCl) with ampicillin. The strains were precultured on Yeast Nitrogen Base (YNB) medium without amino acids and with 5 g/L ammonium sulfate (Sigma-Aldrich Y0626) and 10 g/L glucose. Growth experiments were performed on YNB medium with 1 or 2 % methanol supplemented with 1 g/L yeast extract (Merck 103753). Optical density readings at 600 nm (OD_600_) were used to track growth. For spot assays, swabs from streaked agar plates were pre-cultured twice in 10 mL of 1 x YNB, 1% glucose in sterile 50 mL Falcon tubes. During the exponential phase of the final pre-culture, cells were washed twice in 10 mL of sterile MilliQ water and serially diluted 10-fold up to 10^-4^, prior to spotting 5 μl from each dilution onto the indicated agar plates. Plates were incubated at 30°C for 5 days. Spot assay photos are representative of many repeated experiments. For the Adaptive Laboratory Evolution experiment, three independent lineages of CEN.PK 113-5D with pRS416 where grown on baffled shake flasks with 50 mL YNB medium containing 1 g/L yeast extract or 1 g/L yeast extract with 2 % methanol (48 hours) pulsing with YNB medium with 1 % glucose (24 hours), and incubated at 30°C. OD_600_ measurements were taken before every passage and the cultures were inoculated at a starting OD_600_ of 0.2 if grown on conditional media or OD_600_ of 0.02 if grown on glucose medium.

### DNA extraction and whole-genome sequencing of ALE lineages

Glycerol stocks of the six independent evolutionary lineages were inoculated into 10 mL of YPD medium and grown overnight. Cells were pelleted and washed twice in 10 mL sterile MilliQ water by centrifuging at 4000 x g for 2 mins. Genomic DNA was extracted from pellets using the Yeast DNA Extraction Kit from ThermoFisher (catalog number 78870) according to the instructions. Whole genome sequencing was performed at the Ramaciotti Centre for Genomics using Nextera XT library preparation and NextSeq500 2 x 150 bp sequencing. A minimum of 11 million reads were generated per sample with >95 % of reads having Q20 quality scores, except for methanol evolved lineage B, which had 80 % of reads with at least Q20. An annotated CEN.PK113-7D reference genome was generated by transferring annotated coding sequences with greater than 95 % homology from the S288C reference genome to CEN.PK113-7D FASTA files using Geneious Pro version 11 ^64^. Untrimmed reads were mapped to the annotated CEN.PK113-7D reference genome using Geneious Pro. Each sample had greater than 99 % of reads mapped to the reference with approximately 100-fold coverage. Non-synonymous single nucleotide variants (SNV) within coding sequences were identified at a minimum coverage of 10x, read-variant frequency of 0.9, maximum variant p-value of 10-5, and a minimum strand-bias *p* value of 10^−5^ when exceeding 65% bias. SNVs were annotated onto the consensus sequence of each strain and SNVs present in the methanol-exposed lineages but absent in the yeast extract only lineages were identified as potentially causative mutations.

### Reconstruction of evolved lineage C mutation in *YGR067C*

The methanol-exposed evolutionary lineages had convergent but non-identical mutations in *YGR067C,* which all encoded truncations in the protein through premature stop codons. These mutations were substitution of C to G at nucleotide position 788, deletion of A at position 1877, and substitution of C at position 1176 to G. The C1176G mutation from lineage C was chosen for reverse engineering into the parental CEN.PK113-5D strain due to the proximity of the mutation to a favourable crispr-Cas9 PAM site. The mutation was constructed using a previously described approach whereby a single pRS423 plasmid encoding the Cas9 protein, the guide RNA, and a hygromycin selection marker is PCR amplified using primers that encode a new guide sequence in their 5’ extensions, which also have 20 nt of homology to one another ^65, 66^. This PCR product was co-transformed with mutated *YGR067C* DNA that was PCR amplified from evolved lineage C genomic DNA, with transformants selected for hygromycin resistance on YPD agar plates. Crude genomic DNA was extracted using the lithium acetate-SDS method as previously described ^67^. Transformants were screened by PCR amplifying and Sanger sequencing the putatively mutated region of *YGR067C*. 1 out of 12 tested colonies had the desired mutation using this method. This strain was subsequently referred to as ‘reconstructed EC’.

### ambr^®^ 250 Bioreactor Fermentations

Strains were pre-cultured as above in sterile 50 mL Falcon tubes prior to inoculation at an OD_600_ of 0.02 in ambr**^®^** 250 (Sartorius Stedim) microbial bioreactors with 140 mL of medium. Dissolved oxygen was maintained at a minimum level of 20 % via automatic control of stirring speed, air sparging, and O_2_ sparging. pH was maintained at 5 via the automatic addition of 5 M potassium hydroxide or 1 M phosphoric acid, and temperature was kept at 30 °C. Bioreactors were sampled robotically at indicated time-points, with 1 mL samples stored at -20 °C prior to measurement of growth using OD_600_.

### ^13^C- Methanol Fermentations

Growth experiments in YNB medium supplemented with yeast extract (1 g/L) and ^13^C-methanol (2 %; Sigma-Aldrich 277177) were carried out aerobically (sparging air) at 30°C, agitation of 300 RPM, and with a starting volume of 250 mL of medium using the Multifors 2 bioreactor system (Infors AG). pH was maintained between 4.8 and 5.2 using 3M H_3_PO_4_. For pre-culturing, biological replicates of each strain were grown in liquid YNB medium for approximately 15 h. Cultures were then passaged into a second pre-culture and grown to mid-exponential phase (OD_600nm_ between 1 and 3). A third pre-culture was also grown to mid-exponential phase and washed twice with sterile water before inoculating the experimental cultures at an OD_600_ of 0.02.

### RNA-extraction, sequencing and transcriptome analysis

30 mL of culture was centrifuged at 4 °C 17,000 x g for 10 min, with pellets resuspended in RNA later (Sigma Aldrich R0901) and stored at -20 °C prior to RNA extraction. Total RNA was extracted as previously described ^40^. Library preparation and sequencing was performed at the Ramaciotti Centre for Genomics using a TruSeq Stranded mRNA-seq preparation kit and NextSeq 500 2x75bp sequencing. 30 million reads were generated per sample with Q30 > 93 % and Q20 > 95 %. Untrimmed reads were mapped to the S288C genome with read counting and differential expression analysis carried out using Geneious Pro version 11 ^68^ as previously described ^40^. Approximately 72 million reads were generated per sample. Genes with p-values less than 0.01 were considered differentially expressed. Lists of up and down-regulated genes were analysed for GO term and pathway enrichment using YeastMine at the Saccharomyces Genome Database (https://yeastmine.yeastgenome.org/yeastmine/bag.do). Significant enrichment against the background whole-genome gene list was identified with p-values less than 0.05 and Holm-Bonferroni correction for multiple comparisons. A list of all significant genes that were differentially expressed can be found in Supplementary File 1.

### Analytical methods

#### Off-gas data analysis

Real-time analysis of bioreactor culture off-gas was achieved using a Hiden HPR-20-QIC mass spectrometer (Hiden Analytical) that was connected to the bioreactors. The Faraday Cup detector was used to monitor the intensities of N, Ar, CO_2_, ^13^C-CO_2_, ethanol, ^13^C-ethanol, and ^13^C-methanol at 28, 40, 44, 45, 27, 47, and 30 atomic mass unit (amu), respectively. To increase sensitivity and detect the presence of ^13^C-ethanol from the off-gas data, the Secondary Electron Multiplier (SEM) detector was used to scan any intensities from 15 to 50 amu, with 47 and 48 amu corresponding to ^13^C-ethanol. N intensity (constant during fermentation as nitrogen was an inert gas in our experiments) at 28 amu was used to normalise the intensity from ^13^C-ethanol. The SEM detector scanned the intensities during two to six independent cycles for each bioreactor and at two different time points.

#### Metabolomics

To measure intracellular metabolites samples were taken at 11 hours during the ^13^C-methanol (2 %) fermentations, quenched in methanol and frozen at −80 °C. For extraction, the pellet was resuspended in 2 mLs of 50 % acetonitrile and transferred to 2 mL microcentrifuge tubes with 0.1 mm diameter glass beads, every sample had two technical replicates. Samples were vortexed for 30 s using a Precellys 24 tissue homogenizer (Bertin Instruments) at 30 °C. Three rounds of vortexing were performed allowing the samples to cool completely between rounds. Samples were centrifuged for 3 min at 5,000 x g and the supernatant was transferred to a clean microcentrifuge tube and stored overnight at −80 °C. 2 mL of samples were freeze-dried overnight, then resuspended in 100 μL of water with 10 uM of azidothymidine (AZT) as internal standard, then transferred to HPLC glass inserts for analysis.

Metabolites were analysed using liquid chromatography tandem mass spectrometry (LC-MS/MS) as adapted from ^69–71^. In brief, analyses were performed using a Dionex Ultimate 3000 HPLC system coupled to an ABSciex 4000 QTRAP mass spectrometer. Liquid chromatography was performed using a 50 min gradient, detailed in Supplementary Table 1, with 300 μL/min flowrate, on a Phenomenex Gemini-NX C18 column (150 x 2 mm, 3 μm, 110 A), with a guard column (SecurityGuard Gemini-NX C18, 4 x 2 mm), and column temperature of 55 °C. The mobile phases used were: 7.5 mM aqueous tributylamine (Sigma-Aldrich) with pH adjusted to 4.95 (±0.05) using acetic acid (Labscan) for Solvent A, and acetonitrile (Merck) for Solvent B. Samples were kept at 4 °C in the autosampler and 10 μL were injected for analyses. The HPLC was controlled by Chromeleon 6.80 software (Dionex). Mass spectrometry was achieved using a scheduled multiple reaction monitoring (sMRM) method on the negative ionisation mode. Details of the compound-specific parameters used in the sMRM analysis are listed in Supplementary Table 2. Other hardware parameter values include: ion spray voltage -4500 V, ion source nebuliser (GS1), ion source auxiliary (GS2), curtain (CUR) and collision (CAD) gases were 60, 60, 20 and medium (arbitrary units), respectively, using an in-house ultra-high purity liquid nitrogen tank (BOC). The auxiliary gas temperature was kept at 350 °C. The mass spectrometer was controlled by Analyst 1.6.3 software (AB Sciex). Amounts obtained for each metabolite detected were based on standard curves from serial dilutions of analytical standards, purchase from Sigma, where L1 = 200000, L2 = 100000, L3 = 50000, … L20 = 0.38 nM. This number of standard dilutions guaranteed that the standard curves consisted of at least five data points. Standard mix and pooled samples were regularly injected along the run sequence for quality control. Collected data were processed using MultiQuant 2.1 (AB Sciex).

#### Protein extraction and Mass Spectrometry-based proteomics

Protein samples were centrifuged at 4 °C 17,000 x g for 10 min, the supernatant was discarded and the cell pellet was washed with 1.5 mL of Gibco™ Phosphate-buffered Saline (Life Technologies). Cells were resuspended in lysis buffer (5 % SDS, 50 mM triethylammonium bicarbonate, 100 mM DTT, pH 7.55) and transferred to a microcentrifuge tube containing 0.5 mm diameter glass beads. Cells were disrupted by using a Precellys 24 tissue homogenizer (Bertin Instruments) for 8 rounds of vortexing (30 s vortex, 45 s rest) without cooling. Microcentrifuge tubes were centrifuged for 10 min at 14,000 RPM and supernatant was transferred to a clean microcentrifuge tube and stored at -20 °C. Samples were digested with trypsin following the S-trap™ minis (ProtiFi, LCC.) protocol. The digested peptide mixtures were concentrated using Millipore^®^ ZipTip C18 (Merck) eluting with 70 % acetonitrile. Residual acetonitrile was removed by vacuum centrifugation and peptides resuspended in 5 % acetonitrile, 0.1% formic acid (aqueous) before analysis. Peptides were analysed using a ThermoFisher Scientific UltiMate 3000 RSLCnano UHPLC system. Each sample was initially injected onto a ThermoFisher Acclaim PepMap C_18_ trap reversed-phase column (300 µm x 5 mm nano viper, 5 µm particle size) at a flow rate of 20 µL/min using 2 % acetonitrile (aqueous) for 5 min with the solvent going to waste. The trap column was switched in-line with the separation column (ThermoFisher EasySpray Pepmap RSLC C_18_, 150 µm x 150 mm, 2 µm) and the peptides were eluted using a flowrate of 1.0 µL/min using 0.1 % formic acid in water (buffer A) and 80 % acetonitrile in buffer A (buffer B) as mobile phases for gradient elution. Peptide elution employed a 4-30 % acetonitrile gradient for 40 min followed by 30-50 % acetonitrile for 10 min and 50-95 % acetonitrile for 1 min at 40 °C. The total elution time was 60 min including a 95 % acetonitrile wash followed by re-equilibration. For each sample run, a volume of 2 µL equating to approximately 1 µg of peptide material from protein digestion was injected. The eluted peptides from the C^18^ column were introduced to the MS via a nano-ESI and analysed using the Q-Exactive HF-X. The electrospray voltage was 1.8 kV in positive ion mode, and the ion transfer tube temperature was 275 °C. Employing a top-40ddMS2 acquisition method, full MS-scans were acquired in the Orbitrap mass analyser over the range m/z 350– 1400 with a mass resolution of 120,000 (at m/z 200). The AGC target value was set at 3.00E+06. The 40 most intense peaks with a charge state between 2 and 5 were fragmented in the high energy collision dissociation (HCD) cell with a normalised collision energy (NCE) of 28. MSMS spectra were acquired in the Orbitrap mass analyser with a mass resolution of 15,000 at m/z 200. The AGC target value for MSMS was set to 1.0E+05 while the ion selection threshold was set to 1E+03 counts. The maximum accumulation times were 60 min for full MS-scans and MSMS. For all the experiments, the dynamic exclusion time was set to 25 s, and undetermined charge state species were excluded from MSMS.

Raw files were processed using MaxQuant ^72^ version 1.6.10.43 with the integrated Andromeda search engine ^73^ and the following search parameters: trypsin digestion with a maximum of 2 missed cleavages, carbamidomethyl (C) as a fixed modification and oxidation (M) and acetyl (protein N-term) as variable modifications. The MS data was searched against the SGD protein sequence database (6,630 entries downloaded November 2019). A peptide spectrum match False Discovery Rate (FDR) of 0.01 and a protein FDR of 0.01 were used as protein identification level cut-offs. Label-free quantification (LFQ) ^74^ was performed using MaxQuant LFQ intensities. Protein quantification analysis of the LFQ results was performed using Perseus ^75^ version 1.6.10.0. The data was first filtered to remove identified proteins classified as ‘reverse’, ‘only identified by site’ and ‘potential contaminants’. The two corresponding biological replicates for CEN.PK 113-5D or the reconstructed EC strain were loaded separately and classified as replicates. Proteins that were not present in at least one of the two biological replicates were removed to further trim the dataset. All LFQ values were log2 transformed, the median intensity was subtracted to all intensities for all samples to normalise the distribution, and missing values were imputed to 0. Lastly, a two-sided *t*-test was performed between CEN.PK 113-5D and the reconstructed EC strain, and proteins with FDR adjusted p-values of less than 0.05 were designated as differentially expressed. The log2FC for the significant proteins was then calculated. A list of all significant proteins that were differentially expressed can be found in Supplementary File 2.

## Supporting information

Plasmid maps

## Acknowledgements

This work was funded through the CSIRO Synthetic Biology Future Science Platform, and an internal grant from Macquarie University. The Macquarie University synthetic biology laboratory and The Australian Genome Foundry are funded by Bioplatforms Australia, the New South Wales (NSW) Chief Scientist and Engineer, and the NSW Government’s Department of Primary Industries. There was no funding support from the European Union for the experimental part of the study. However, K.V. acknowledges support also from the European Union’s Horizon 2020 research and innovation programme under grant agreement N810755. M.I.E and T.C.W thank E.M for his guidance and contribution whilst designing the ^13^C-labelling bioreactor experiments. M.I.E thanks Bradley W. Wright from Macquarie University for his guidance during the proteomics analysis.

## Author contributions

M.I.E and T.C.W conceived the study and designed the experiments. M.I.E performed the experiments, with contributions from T.C.W. M.I.E and T.C.W analysed data. M.I.E, E.M, K.V and R.A.G.G designed and performed the ^13^C-labelling and bioreactor experiments. M.P designed the method to detect intracellular ^13^C-labelled metabolites. C.S, T.C.W, I.S.P and I.T.P participated in the design, support, and coordination of the project. All authors read and approved the final manuscript.

## Materials and correspondence

Thomas C. Williams, Ian T. Paulsen

