## Supplementary Information for "Engineering and Evolution of Methanol Assimilation in *Saccharomyces cerevisiae*"

**Supplementary Fig. 1. Spot-assays on different carbon sources of *S. cerevisiae* strains expressing different methanol assimilation pathways.** Growth of serially 10-fold diluted strains on solid YNB medium with indicated carbon sources. The empty vector control is highlighted in black and the yeast XuMP strain is highlighted in red. The ‘hybrid’ XuMP strain and variations thereof are in green. Lastly, the bacterial RuMP strain and variations thereof are shown in blue. Yeast Nitrogen Base (YNB), Yeast Extract (YE), Methanol (MeOH). Images were taken after incubating at 30 ºC for 5 days. The Yeast XuMP assays are from different plates but following the same methodology and with an empty vector control.

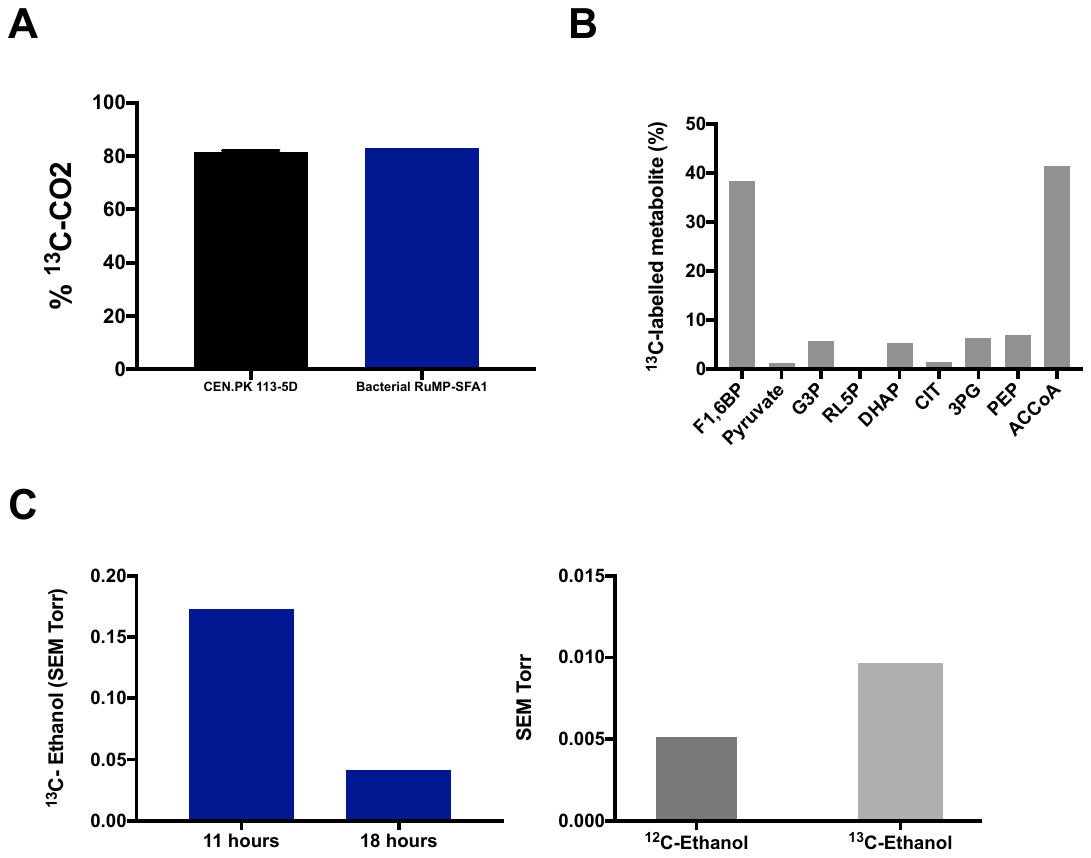

**Supplementary Fig. 2. ^13^C-methanol fermentations of the bacterial RuMP*-pTDH3-SFA1* strain.** **A.** Percentage of ^13^C-CO_2_/CO_2_ production of the empty vector control or bacterial RuMP-*pTDH3-SFA1* strains on yeast extract ^13^C-methanol (2 %) medium **B.** Percentage of ^13^C-labelled intracellular metabolites in the bacterial RuMP-*pTDH3-SFA1* strain, metabolites are universally (fully) labelled with ^13^C. F1,6BP, fructose 1,6-bisphosphate; G3P, glyceraldehyde-3-phosphate; RL5P, ribulose-5-phosphate; DHAP, dihydroxyacetone phosphate; CIT, citrate; 3PG, 3-phosphoglyceric acid; PEP, phosphoenolpyruvate; ACCoA, acetyl-coenzyme A. Data points represent one biological replicate. **C.** ^13^C-ethanol was produced the bacterial RuMP-*pTDH3-SFA1* strain. The signal intensity was normalised to the inert gas nitrogen, and then to biomass. Data shows the average ^13^C-ethanol intensity at 47 amu for one biological replicate during independent scanning cycles using a Hiden HPR-20-QIC mass spectrometer. Ratio of ^12^C-ethanol and ^13^C-ethanol produced by the bacterial RuMP-*pTDH3-SFA1* strain at 18 hours. The signal intensity was normalised to the inert gas nitrogen, and then to biomass. Data shows the average ^12^C-ethanol and ^13^C-ethanol intensity at 31 and 33 amu, respectively for one biological replicate during independent scanning cycles using a Hiden HPR-20-QIC mass spectrometer. SEM, Secondary Electron Multiplier; amu, atomic mass unit.

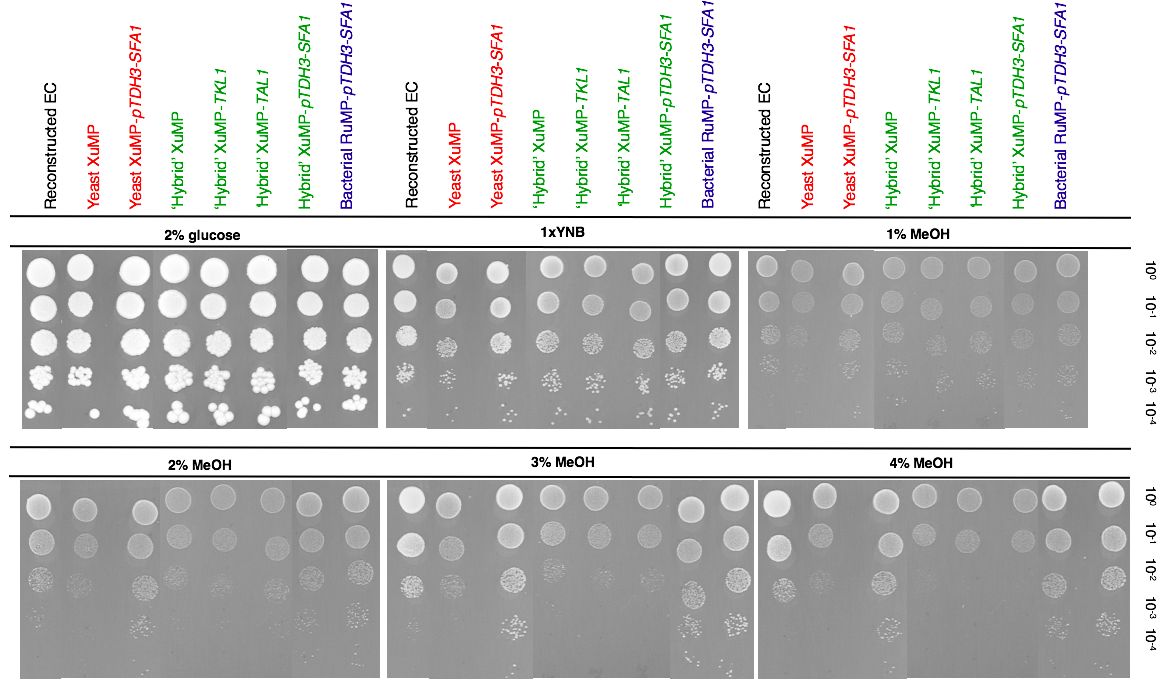

**A**

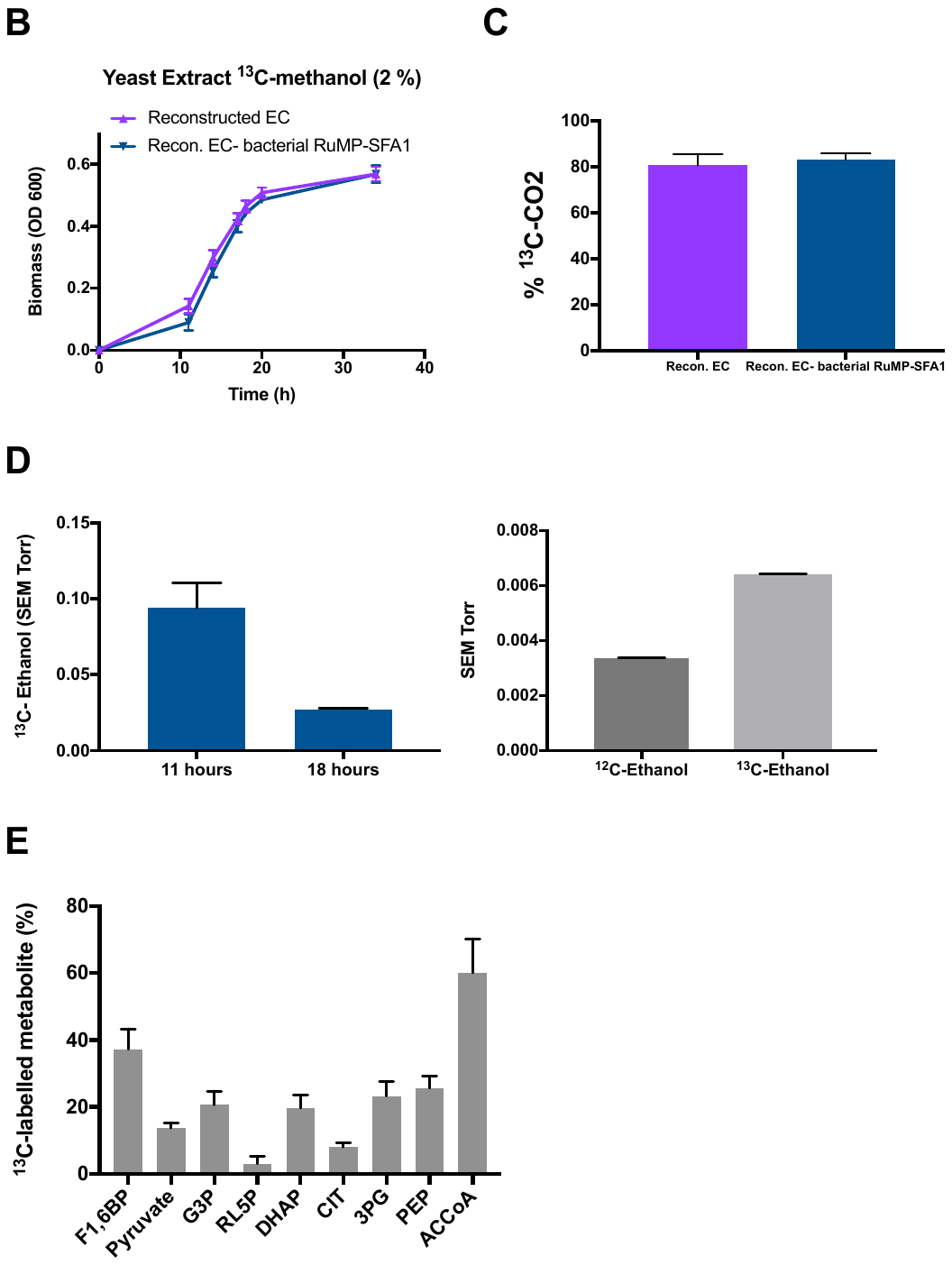

**Supplementary Fig. 3. Reconstructed evolved strain with engineered methanol assimilation pathways. A.** Growth on solid 1x Yeast Nitrogen Base medium with different carbon sources was tested using serial 10-fold dilutions of the reconstructed evolved strain with an empty vector or with the engineered methanol assimilation pathways. Yeast Nitrogen Base (YNB), Yeast Extract (YE), Methanol (MeOH). Images were taken after incubating at 30 ºC for 6 days. **B.** Growth profile of the reconstructed evolved strain with an empty vector and with the bacterial RuMP-*pTDH3-SFA1* pathway, strains were grown in liquid YNB medium with 2 % ^13^C-methanol supplemented with 0.1 % yeast extract. **C.** Percentage of ^13^C-CO_2_/CO_2_ production of the reconstructed evolved strain with an empty vector or with the bacterial RuMP-*pTDH3-SFA1* pathway on yeast extract ^13^C-methanol (2 %) medium **D.** ^13^C-ethanol was produced by the reconstructed evolved strain with the bacterial RuMP-*pTDH3-SFA1* pathway. The signal intensity was normalised to the inert gas nitrogen, and then to biomass for each strain. Data shows the average ^13^C-ethanol intensity at 47 amu for two biological replicates during independent scanning cycles using a Hiden HPR-20-QIC mass spectrometer. Error bars are the standard deviation of the ^13^C-ethanol intensity. Ratio of ^12^C-ethanol and ^13^C-ethanol produced by the reconstructed evolved strain with the bacterial RuMP-*pTDH3-SFA1* pathway at 18 hours. The signal intensity was normalised to the inert gas nitrogen, and then to biomass for each strain. Data shows the average ^12^C-ethanol and ^13^C-ethanol intensity at 31 and 33 amu, respectively for two biological replicates during independent scanning cycles using a Hiden HPR-20-QIC mass spectrometer. Error bars are the standard deviation of the ^12^C-ethanol or ^13^C-ethanol intensities. SEM, Secondary Electron Multiplier; amu, atomic mass unit. **E.** Percentage of ^13^C-labelled intracellular metabolites in the reconstructed evolved strain with the bacterial RuMP-*pTDH3-SFA1* pathway, metabolites are universally (fully) labelled with ^13^C. F1,6BP, fructose 1,6-bisphosphate; G3P, glyceraldehyde-3-phosphate; RL5P, ribulose-5-phosphate; DHAP, dihydroxyacetone phosphate; CIT, citrate; 3PG, 3-phosphoglyceric acid; PEP, phosphoenolpyruvate; ACCoA, acetyl-coenzyme A. Data points represent the average of two biological replicates and error bars are the standard deviation.

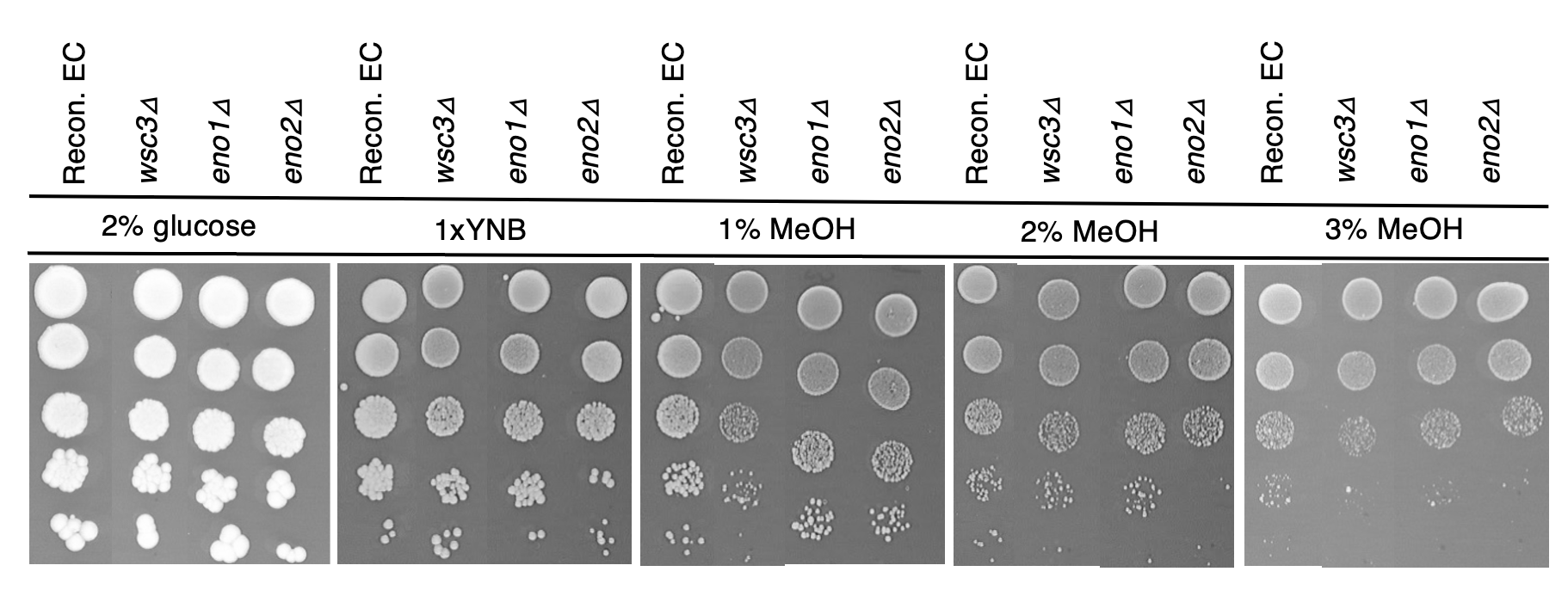

**Supplementary Fig. 4. Growth in methanol of different gene deletions to test their putative involvement *in S. cerevisiae*’s native methanol assimilation.** Growth on solid 1x Yeast Nitrogen Base medium with different carbon sources was tested using serial 10-fold dilutions of the reconstructed evolved strain with an empty vector or *WSC3*, *ENO1* or *ENO2* deletions. Yeast Nitrogen Base (YNB), Yeast Extract (YE), Methanol (MeOH). Images were taken after incubating at

**Supplementary Table 1.** Liquid chromatography gradient profile

| **Time (min)** | **% B** |
| --- | --- |
| 0 | 0 |
| 8 | 0 |
| 20 | 20 |
| 30 | 27 |
| 31 | 100 |
| 33 | 100 |
| 34 | 0 |
| 50 | 0 |

**Supplementary Table 2**. Metabolite-specific parameters used in the acquisition of the sMRM data.

| **Metabolite** | **Q1 (m/z)** | **Q3 (m/z)** | **RT (min)** | **DP (V)** | **EP (V)** | **CE (V)** | **CXP (V)** |
| --- | --- | --- | --- | --- | --- | --- | --- |
| PYR | 87.02 | 43.0 | 12.0 | -45 | -10 | -12 | -1 |
| PYR_U13C | 90.00 | 45.0 | 12.0 | -45 | -10 | -12 | -1 |
| LAC | 88.95 | 42.9 | 8.4 | -45 | -10 | -18 | -5 |
| LAC_U13C | 92.00 | 45.0 | 8.4 | -45 | -10 | -18 | -5 |
| FUM | 115.01 | 70.9 | 20.8 | -45 | -10 | -12 | -1 |
| FUM_U13C | 119.00 | 74.0 | 20.8 | -45 | -10 | -12 | -1 |
| SUC | 117.01 | 73.0 | 18.4 | -45 | -10 | -16 | -3 |
| SUC_U13C | 121.00 | 76.0 | 18.4 | -45 | -10 | -16 | -3 |
| OAA | 130.93 | 86.9 | 20.0 | -25 | -10 | -10 | -5 |
| OAA_U13C | 135.00 | 90.0 | 20.0 | -25 | -10 | -10 | -5 |
| MAL | 133.00 | 70.8 | 19.4 | -40 | -10 | -22 | -3 |
| MAL_U13C | 137.00 | 74.0 | 19.4 | -40 | -10 | -22 | -3 |
| KGA | 144.95 | 100.8 | 20.1 | -40 | -10 | -12 | -5 |
| KGA_U13C | 150.00 | 105.0 | 20.1 | -40 | -10 | -12 | -5 |
| PEP | 166.83 | 79.0 | 21.7 | -40 | -10 | -18 | -5 |
| PEP_U13C | 170.00 | 79.0 | 21.7 | -40 | -10 | -18 | -6 |
| GA3P | 168.84 | 97.0 | 11.8 | -40 | -10 | -10 | -5 |
| GA3P_U13C | 172.00 | 97.0 | 11.8 | -40 | -10 | -10 | -5 |
| DHAP | 168.84 | 97.0 | 11.9 | -50 | -10 | -14 | -5 |
| DHAP_U13C | 172.00 | 97.0 | 11.9 | -50 | -10 | -14 | -5 |
| 3PG | 184.91 | 97.0 | 21.1 | -50 | -10 | -20 | -5 |
| 3PG_U13C | 188.00 | 97.0 | 21.1 | -50 | -10 | -20 | -5 |
| CIT | 190.96 | 110.9 | 21.8 | -50 | -10 | -18 | -7 |
| CIT_U13C | 197.00 | 116.0 | 21.8 | -50 | -10 | -18 | -7 |
| R5P | 228.94 | 96.9 | 8.7 | -20 | -10 | -30 | -15 |
| R5P_U13C | 234.00 | 96.9 | 8.7 | -20 | -10 | -30 | -15 |
| RL5P | 228.92 | 96.9 | 11.0 | -20 | -10 | -30 | -15 |
| RL5P_U13C | 234.00 | 96.9 | 11.0 | -20 | -10 | -30 | -15 |
| G1P | 259.02 | 78.8 | 9.8 | -20 | -10 | -30 | -15 |
| G1P_U13C | 265.00 | 78.8 | 9.8 | -20 | -10 | -30 | -15 |
| G6P | 258.89 | 96.7 | 8.0 | -20 | -10 | -30 | -15 |
| G6P_U13C | 265.00 | 96.7 | 8.0 | -20 | -10 | -30 | -15 |
| F6P | 259.02 | 96.8 | 9.1 | -20 | -10 | -30 | -15 |
| F6P_U13C | 265.00 | 96.8 | 9.1 | -20 | -10 | -30 | -15 |
| F16BP | 339.08 | 96.9 | 21.4 | -20 | -10 | -30 | -15 |
| F16BP_U13C | 345.00 | 96.9 | 21.4 | -20 | -10 | -30 | -15 |
| GLYOX | 73.00 | 45.0 | 5.7 | -45 | -10 | -12 | -1 |
| GLYOX_U13C | 75.00 | 46.0 | 5.7 | -45 | -10 | -12 | -1 |
| GLYCO | 75.00 | 47.0 | 5.8 | -35 | -10 | -14 | -3 |
| GLYCO_U13C | 77.00 | 48.0 | 5.8 | -35 | -10 | -14 | -3 |
| UDPG | 565.18 | 323.0 | 20.8 | -90 | -10 | -34 | -7 |
| UDPG_U13C | 571.00 | 323.0 | 20.8 | -90 | -10 | -34 | -7 |
| UDPGA | 579.14 | 79.1 | 28.6 | -90 | -10 | -108 | -1 |
| UDPGA_U13C | 585.00 | 79.1 | 28.6 | -90 | -10 | -108 | -1 |
| ACOA | 808.17 | 79.1 | 32.2 | -125 | -10 | -54 | -5 |
| ACOA_U13C | 810.00 | 79.1 | 32.2 | -125 | -10 | -54 | -5 |
| UDPNAc | 605.86 | 78.7 | 20.9 | -95 | -10 | -106 | -1 |
| UDPNAc_U13C | 612.00 | 78.7 | 20.9 | -95 | -10 | -106 | -1 |
| ACO | 172.94 | 84.9 | 22.0 | -30 | -10 | -18 | -5 |
| ACO_U13C | 177.00 | 87.0 | 22.0 | -125 | -10 | -54 | -5 |
| CMP | 322.07 | 78.8 | 11.9 | -65 | -10 | -66 | -3 |
| UMP | 323.01 | 78.8 | 13.4 | -60 | -10 | -64 | -3 |
| AMP | 346.02 | 78.6 | 15.9 | -70 | -10 | -62 | -3 |
| GMP | 362.05 | 78.9 | 14.1 | -60 | -10 | -62 | -3 |
| UDP | 403.03 | 78.8 | 21.5 | -60 | -10 | -74 | -3 |
| ADP | 426.07 | 78.8 | 22.0 | -85 | -10 | -74 | -3 |
| GDP | 442.06 | 78.9 | 21.3 | -70 | -10 | -76 | -3 |
| CTP | 481.95 | 158.6 | 27.8 | -75 | -10 | -36 | -11 |
| UTP | 483.06 | 158.8 | 29.1 | -65 | -10 | -42 | -7 |
| ATP | 506.10 | 158.7 | 29.4 | -85 | -10 | -40 | -11 |
| GTP | 522.00 | 158.7 | 28.8 | -80 | -10 | -42 | -11 |
| NAD | 662.25 | 540.0 | 13.3 | -50 | -10 | -20 | -9 |
| NADH | 664.20 | 78.8 | 22.3 | -110 | -10 | -98 | -1 |
| NADP | 742.20 | 620.0 | 21.2 | -45 | -10 | -24 | -11 |
| NADPH | 744.10 | 79.1 | 29.2 | -120 | -10 | -116 | -1 |
| Cre-P | 209.74 | 78.8 | 19.0 | -35 | -10 | -16 | -3 |
| AZT | 265.80 | 223.0 | 14.0 | -70 | -10 | -16 | -1 |

***Note:*** U13C: denotes ^13^C universally labelled metabolite, Q, quadrupole; RT, retention time; DP, declustering potential; EP, entrance potential; CE, collision energy; CXP, collision cell exit potential; V, volts.
